# Connecting the Dots: PHF13 and cohesin promote polymer-polymer phase separation of chromatin into chromosomes

**DOI:** 10.1101/2022.03.04.482956

**Authors:** Francesca Rossi, Rene Buschow, Laura V. Glaser, Tobias Schubert, Hannah Staege, Astrid Grimme, Hans Will, Thorsten Milke, Martin Vingron, Andrea M. Chiariello, Sarah Kinkley

## Abstract

How interphase chromatin compacts into mitotic chromosomes has eluded researchers for over a century. Here we show that PHF13, a tightly regulated H3K4me epigenetic reader, and cohesin, a mediator of chromatin architecture, cooperate to drive polymer-polymer-phase-separation (PPPS) and higher order compaction of chromatin into chromosomes. PHF13 interacts with cohesin, shows similar dynamics during mitosis, and their co-depletion dramatically impairs mitotic condensation. Mechanistically, we demonstrate that PHF13 stabilizes cohesin chromatin interactions and that itself oligomerizes, resulting in a polymer with increased chromatin avidity and the ability to bridge neighboring and distant nucleosomes. Consistently, molecular dynamic simulations modelling the ability of PHF13 and cohesin to drive chromatin phase separations recapitulated our in vivo observations and are in line with 2-step condensation models.

## Introduction

At the transition from interphase to mitosis, the hierarchical organization of chromatin is massively restructured resulting in individual rod-shaped chromosomes, which are several fold more compacted than interphase chromatin. How this is orchestrated remains poorly understood and has been fascinating research since its initial report by Walther Fleming in 1878. Early models of mitotic higher chromatin order formation postulated that chromatin fibers are successively folded in a hierarchical regular or irregular manner upon themselves to form compacted metaphase chromosomes (Belmont et al., 1987; Sedat and Manuelidis, 1978). However, computational simulations using only the successive folding of the chromatin fiber upon itself end up forming spherical rather than rod-like structures (Marko and Siggia, 1997), implying inaccuracies in this model. This lead to the hierarchical folding and axial glue (Kireeva et al., 2004) and the polymer melt (Naumova et al., 2013) models, which propose that chromosome condensation is a two-step cascade, initially involving interactions between neighboring nucleosomes which locally fold to linearly compact the fiber, followed by longer distant nucleosome interactions, to promote axial compaction. Consistently, Hi-C contact data derived from synchronized interphase and mitotic cells supported these models and proposed that chromosome condensation and shaping should be viewed as two distinct and successive processes (Gibcus et al., 2018; Green et al., 2012). In the last two decades, significant advances have been made in terms of our understanding of mitotic chromosome shaping (Gibcus et al., 2018; Samejima et al., 2018), while we still do not understand the early mechanisms mediating mitotic compaction.

Novel insights into how mitotic condensation may be orchestrated have been emerging from a plethora of phase separation studies that support liquid-liquid phase separation (LLPS), gel-like phase separation (GLPS) and polymer-polymer phase separation (PPPS) playing a central role in genome organization and compaction (Gibson et al., 2019; Kantidze and Razin, 2020; Ryu et al., 2021). Chromatin architectural changes mediated by phase separation require multivalent protein nucleic acid interactions and saturating concentrations of the effector that localize at nucleation sites, making the reaction energetically favorable to enforce their growth and stability (Banani et al., 2017; Erdel and Rippe, 2018). Several studies have demonstrated the ability of H1, HP1α, CBX proteins and other cationic proteins to promote phase separation in the presence of DNA or chromatin, driving compaction and heterochromatinization (DeRouchey et al., 2013; Grau et al., 2011; Larson et al., 2017; Shakya et al., 2020; Strom et al., 2017). Adding to this list, Cees Deeker and colleagues have recently found that the cohesin holocomplex is capable of polymer-polymer phase separation, initiated by DNA loops that nucleate long distant bridging, termed bridging induced phase separation (Ryu et al., 2021), consistent with earlier molecular dynamic simulations (Brackley et al., 2013; Conte et al., 2020). It remains however to be experimentally demonstrated whether mitotic chromosome formation is driven by novel mitotic phase separation processes.

Cohesin is a SMC containing complex composed of SMC1, SMC3, RAD21 and SA1 or SA2, that is conserved from yeast to mammals. Cohesin is bound to chromatin throughout the cell cycle and orchestrates chromatin architecture in a variety of processes (Banning et al., 2010; Ibrahim and Mundlos, 2020; Nasmyth and Haering, 2009; Rao et al., 2017; Unal et al., 2007) including chromosome condensation (Heidinger-Pauli et al., 2010; Tedeschi et al., 2013). Post replication cohesin forms dimer complexes (Holzmann et al., 2019; Shi et al., 2020) increasing its chromatin affinity until metaphase arguing that increased cohesin chromatin levels may be important for mitotic condensation. Consistently, cohesin depletion compromises chromatin condensation, results in loss of mitotic loops and increases mitotic interchromatin interactions whereas its increased chromatin association promotes chromatin condensation in both yeast and mammals (Costantino et al., 2020; Heidinger-Pauli et al., 2010; Lopez-Serra et al., 2013; Tedeschi et al., 2013). PHF13 is a H3K4 methyl epigenetic reader protein that is also capable of modulating chromatin architecture in a variety of processes (Bordlein et al., 2011; Chung et al., 2016; Kinkley et al., 2009; Mund et al., 2012). Similar to cohesin, PHF13 is bound to chromatin throughout the cell cycle, its chromatin association increases at the G2/M transition and it has been implicated in mitotic chromosome compaction (Kinkley et al., 2009). Here we present data that PHF13 and cohesin cooperate to promote polymer-polymer phase separation (PPPS) of interphase chromatin to initiate higher order chromosome condensation. Our findings are consistent with two-step condensation reactions where PHF13 oligomerizes promoting linear compaction of chromatin via polymer induced phase separation (PIPS) and where cohesin subsequently promotes bridging induced phase separation (BIPS) through long distant interactions. Together, our work identifies PHF13, cohesin and PPPS as driving factors of mitotic chromosome condensation, adding important insights to a ~150-year old riddle.

## Results

### PHF13 and cohesin interact and co-localize in mitosis

PHF13 is a conserved (Figure S1A) H3K4me3 epigenetic reader and regulator of chromatin architecture (Chung et al., 2016; Kinkley et al., 2009; Mund et al., 2012). It is composed of several regulatory domains (Figure 1A and S1A), including nuclear (NLS) and nucleolar (NoLS; Figure S1B) localizing sequences, histone and DNA binding domains (PHD and polycationic stretch embedded in the NLS, respectively), two PEST domains which regulate its half-life and a conserved N-terminal domain (NTD) of unknown function (Chung et al., 2016; Kinkley et al., 2009).

**Figure 1.**
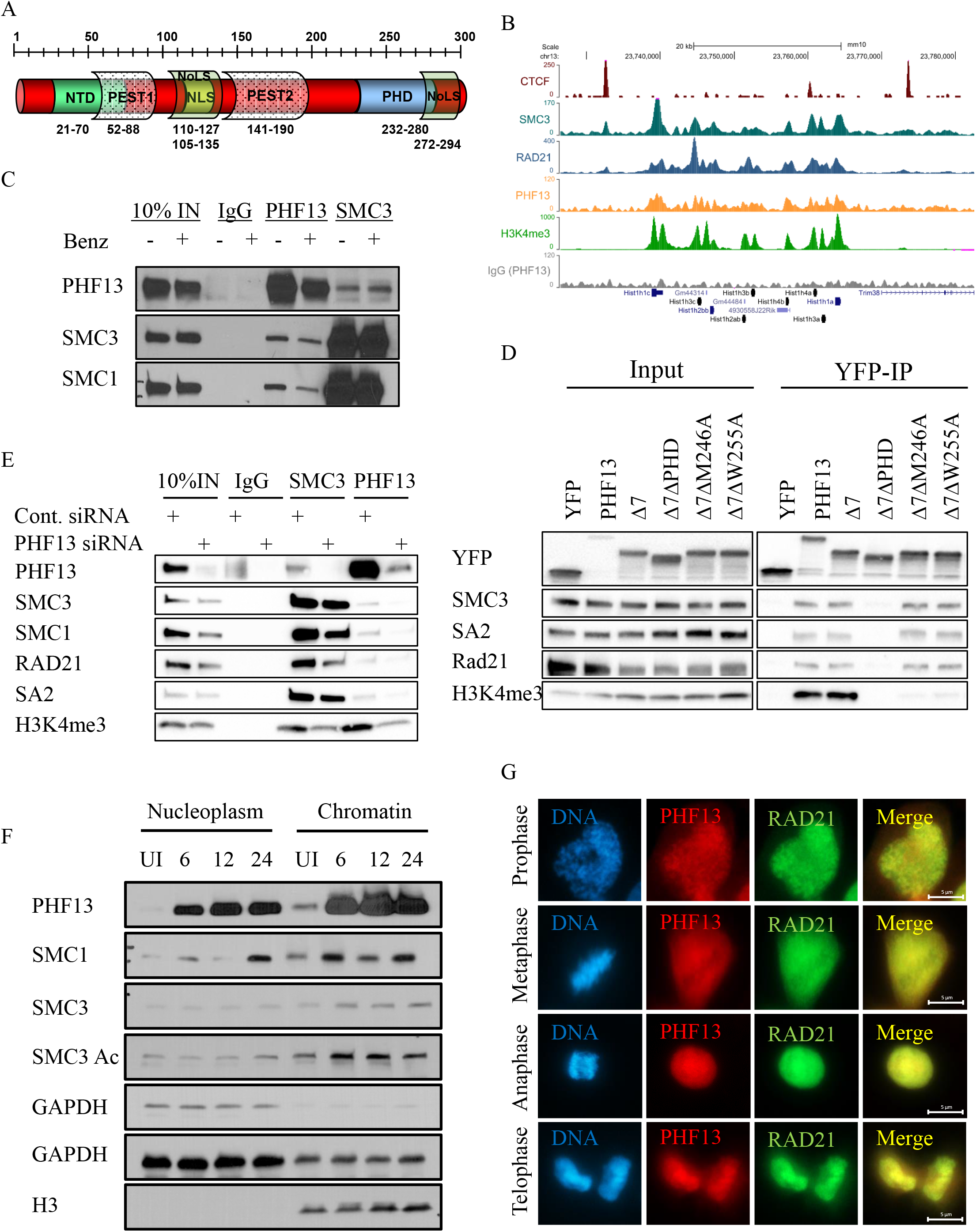
PHF13 and cohesin interact. (A) Schematic depiction of PHF13 domain structure. (B) UCSC browser snapshot: Overlap of RPKM normalized ChIPseq signals at several genomic loci. (C-E) IP:WB; Endogenous IP of PHF13 and SMC3 from U2OS chromatin lysate treated with (+) or without (-) Benzonase and WB for PHF13, SMC3 and SMC1 (C). (D) GFP-TRAP from U2OS cells expressing YFP-tagged proteins. WB for YFP, PHF13, SMC3, SA2 and H3K4me3. (E) Endogenous IP of SMC3 and PHF13 from wt (cont. siRNA) or PHF13 depleted (36h) lysates. WB for SMC3, SMC1, RAD21, SA2 and H3K4me3. (F) WB of fractionated nucleoplasmic and chromatin lysate from uninduced (UI) or induced (6, 12, 24h) U2OS PHF13 clone 5 and probed for PHF13, SMC1, SMC3, SMC3Ac, GAPDH and H3. (G) IF images of mitotic cells co-stained for PHF13 and RAD21. DNA was stained with Hoechst. Images were taken at 100x magnification and captured on a Zeiss Axio Observer. PHF13 was precipitated with a rabbit peptide Ab S173 and detected in WB and IF using rat mAb 6F6 (C, E and G).

PHF13 was previously implicated in modulating mitotic chromatin architecture (Kinkley et al., 2009) motivating us to explore its role in this process and to elucidate the mechanisms involved. As a starting point, we were intrigued by a putative interaction of PHF13 with cohesin detected in an earlier mass spectrometry study (data not shown). In an effort to gather more evidence in support of a relationship with cohesin, we analyzed publicly available ChIP sequencing data sets for PHF13 and cohesin (SMC3 and RAD21), to see if they have a genome-wide overlap (Figure 1B and S1C and D). This analysis demonstrated that PHF13 and cohesin do share a common genomic footprint, at H3K4me3 demarcated landscapes (Figure 1B and S1C and D), underlining a spatial genomic correlation between them. To obtain biochemical evidence for an interaction between PHF13 and cohesin we looked for their ability to interact in endogenous reciprocal co-immunoprecipitation experiments (Figure 1C). These experiments validated that both PHF13 and SMC3 were able to co-precipitate one another as well as SMC1 in both the absence and presence of Benzonase, implying that the interaction is not mediated by RNA or DNA (Figure 1C) and that they interact in sub-stoichiometric ratios. To demonstrate specificity, we mapped the interaction using YFP-tagged PHF13 mutant proteins (Figure 1D) and we reciprocally immunoprecipitated endogenous PHF13 and cohesin (SMC3) from wild-type and PHF13 depleted chromatin lysate (Figure 1E). Cohesin mapped to the last half of PHF13 (Δ7; 150-300) and more precisely to PHF13’s PHD domain in nuclease digested lysates (Figure 1D). Importantly, point mutations (M246A and W255A) in the PHD domain which perturb PHF13 binding to H3K4me3 (Chung et al., 2016), do not disrupt PHF13’s interaction with cohesin, indicating that the interaction is independent of a chromatin substrate (Figure 1C and D) and identifying a novel interaction of PHF13’s PHD domain. Furthermore, PHF13 depletion reduced its precipitation in PHF13 and SMC3 immunoprecipitations and abrogated the co-precipitation of cohesin proteins with PHF13 (Figure 1E) confirming the specificity of these interactions. Interestingly, SMC3 showed reduced precipitation of H3K4me3 in PHF13 depleted lysates, suggesting that PHF13 may be involved in cohesin chromatin recruitment or stability (Figure 1E). To test this further, we induced PHF13 expression in a doxycycline inducible cell line and looked by immunoblotting of fractionated lysates for the impact on cohesin nucleoplasm and chromatin levels (Figure 1F). Induction of PHF13 expression (6h, 12h and 24h) lead to increased cohesin (SMC1 and SMC3) and acetylated SMC3 chromatin levels (Figure 1F), a modification that stabilizes cohesin chromatin association (Beckouet et al., 2016) supporting a correlation between PHF13 and cohesin chromatin levels. Finally, to obtain evidence for a mitotic relationship, we performed indirect immunofluorescence against endogenous PHF13 and RAD21 and looked for their co-localization through mitosis (Figure 1G). This revealed that PHF13 and RAD21 show a similar biphasic chromatin localization during mitosis, namely chromatin associated during prophase and late anaphase/telophase and predominantly cyto-/nucleoplasmic during metaphase/anaphase, consistent with a spatial temporal correlation between PHF13 and cohesin during mitosis. Together, these findings demonstrate that PHF13 and cohesin interact, share a genome-wide profile at H3K4me3 demarcated landscapes, that PHF13 influences cohesin chromatin association and that they show a spatial temporal correlation during mitotic transition and progression, supporting the possibility of a functional relationship.

### PHF13 promotes higher chromatin order

The observation that PHF13 influences cohesin chromatin association led us to question if PHF13 concentration correlates with chromatin compaction. To address this question in an unbiased manner, we determined the cell cycle distribution of asynchronous cells using high throughput microscopy and calculated the spearman correlation coefficient between PHF13 concentration and DNA compaction in different cell cycle states (Figure 2A). Using SPY650 DNA dye and the parameters of sum DNA intensity and standard deviation (SD) of DNA intensity, we could differentiate cells with 2N, 3N and 4N and further distinguish mitosis from G2 (Figure 2A). Supporting the accuracy of the cell cycle distribution, H3S10 phosphorylation exclusively overlapped with cells that were called mitotic and not G2 (Figure S2A and S2B). We next estimated the total (standard fixation) and chromatin bound (pre-extraction) PHF13 levels throughout the cell cycle, confirming that PHF13 is predominantly chromatin bound and that it increases at mitotic transition (Figure 2A) as previously described (Kinkley et al., 2009). Using Spearman correlations (ρ), we evaluated the relationship between chromatin bound PHF13 and the SD of DNA intensity, a principle component which separates interphase and mitotic DNA states (Figure 2A and S2C). This revealed that PHF13 chromatin concentration showed the strongest correlation with early mitosis (Figure2A and S2C), consistent with a possible role of PHF13 in mitotic compaction under physiological conditions.

**Figure 2.**
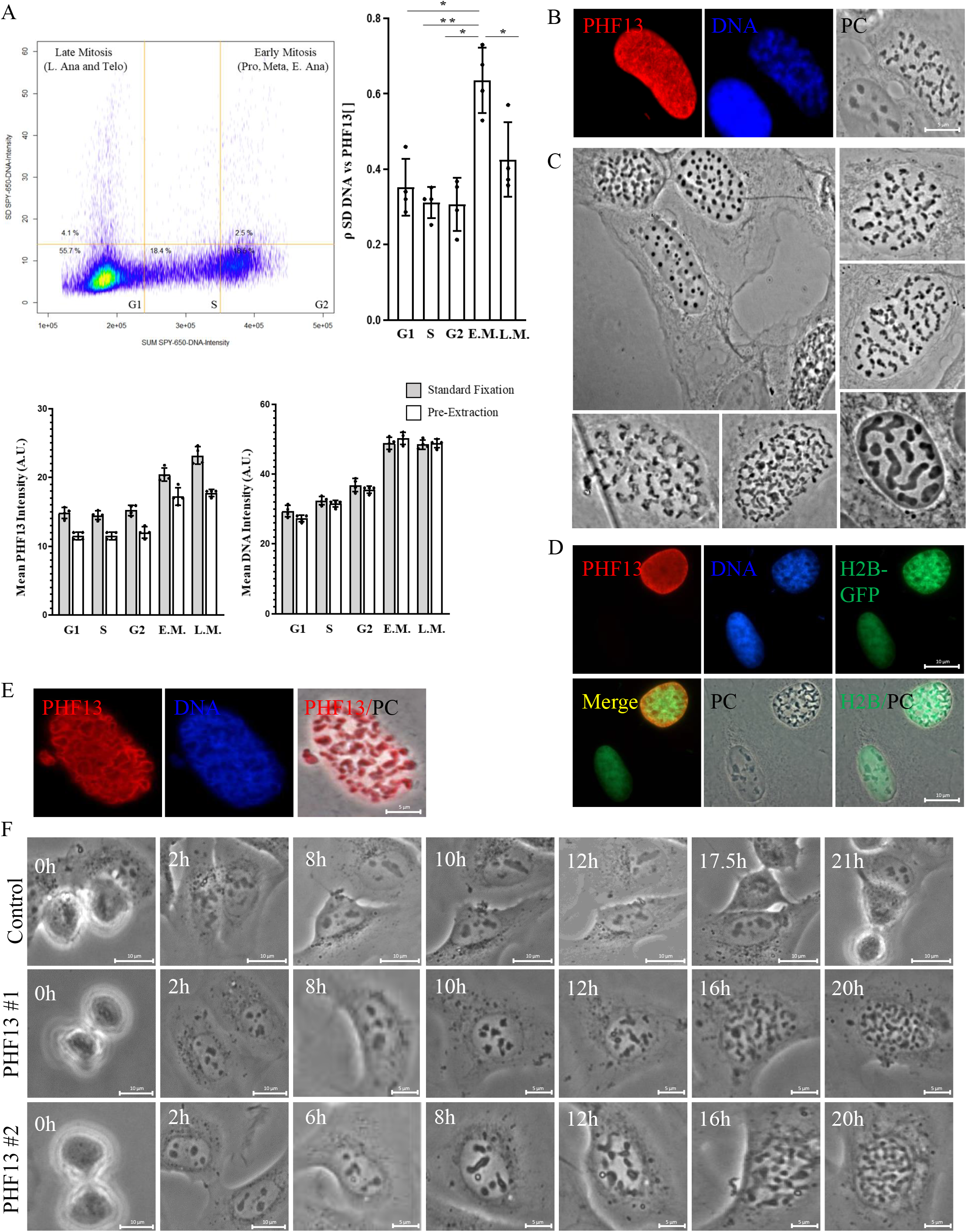
PHF13 promotes higher chromatin order. (A) Spearman correlation of PHF13 concentration in relation to chromatin state was measured in asynchronous cells using the sum and standard deviation of SPY650 intensity to segregate cells into individual cell cycle phases and compaction states, in relation to PHF13 mean signal intensity. (B-E) Indirect IF of U2OS (B and E) or U2OS H2B-GFP (D) cells transfected with 1 μg PHF13 for 24h and paraformaldehyde fixed (B,C and D) or pre-extracted prior to fixation (E). PHF13 was detected with rabbit peptide Ab CR53 (B,E) or rat mAb 6F6 (D) and DNA was stained with Dapi. (F) Live-cell images of control or PHF13 transfected U2OS cells. Images were captured on a Zeiss Cell Discoverer (A) and Axiophot Phase Fluorescence microscope (B-E) or on a Nikon BioStation live cell microscopy (F).

To better observe the impact of PHF13 expression on chromatin state, we transiently expressed PHF13 in U2OS cells and looked at them by indirect immunofluorescence and phase contrast microscopy (Figure 2B–F). Strikingly, PHF13 over expression coincided with global chromosome condensation with a notable refractive difference from the nucleoplasm visible in phase contrast (PC) and Hoechst (DNA) imaging (Figure 2B–C). This phenotype, has only been previously described for cohesin (Tedeschi et al., 2013), and confirms PHF13’s ability to promote higher chromatin order states. We repeated these experiments in U2OS cells stably expressing H2B-GFP and found consistently that H2B-GFP reorganized with the phase contrast compacted structures (Figure 2D). Interestingly, pre-extraction of soluble nuclear proteins prior to fixation revealed that chromatin bound PHF13 forms filaments that encapsulate the compacted chromatin structures (Figure 2E and S2D) suggesting that PHF13 oligomerizes concomitantly with global chromosome condensation and implicating a polymer-induced phase separation (PIPS) mechanism. To understand the distinct chromatin morphologies as well as the dynamics, we performed live cell imaging of U2OS cells transiently expressing PHF13 (Figure 2F, S2E and supplementary movies). To calibrate our approach we first expressed GFP-PHF13 and followed in real-time to determine the kinetics of PHF13 expression post-transfection. GFP was consistently detected shortly after cells completed mitosis, reflecting uptake of plasmid DNA following mitotic nuclear envelope breakdown and expression in G1 phase (data not shown). Based on these observations, we chose telophase as our time point zero to evaluate the kinetics of condensation in our non-tagged PHF13 live cell studies (Figure 2F, S2E and supplementary videos). In control and non-transfected cells the time from mitosis 1 (telophase) to mitosis 2 (telophase) ranged from 17h-22h (Figure 2F-upper panel). Based on this and knowledge from synchronization experiments, we estimated that G1 occurs in U2OS cells from 0-6h post telophase, S-phase from 6-10h post telophase, G2 from 10h-16h post telophase and M-phase from 17h + post telophase, with cell cycle delays prolonging this cycle. With this in mind, we found that between 4 and 10 hours post-telophase that the nucleoli of PHF13 expressing cells started to darken and appeared to nucleate the compacting chromatin (Figure 2F, S2E and supplementary movies). Interestingly, in cells where the nucleolar darkening and nucleation began early, i.e. 4-6 hours post-telophase (Figure S2E; red arrows, PHF13 movie 1: left cell and PHF13 movie 4: left cells) a time point reflecting G1 in U2OS cells, the chromatin compacted into large irregular globular structures by 8-10h. In contrast, in cells where nucleation initiated later i.e. 8-10 hours post telophase, a time point more consistent with S-phase (Figure 2F and S2E; blue arrows and cell movie 1: right cell, cell movie 2: both cells and cell movie 4: right cells), the chromatin ultimately compacted into prophase-like chromosome morphologies with unresolved sister chromatids by 12-16h. In addition, there were a few rare exceptions to these cases where the chromatin formed regular spherical condensates (Figure S2E; PHF13#5 and cell movie 5). In these cases, the nucleolar darkening was observed around 7h post-telophase (late G1/early S-phase) and the spherical condensates formed much later at around 21h and lasted for extended periods of time (>32h post telophase), suggesting that these cells encounter a cell cycle block and that the spherical chromatin condensates reflect individual chromosomes compacted into collapsed globules. In all cases, the compacted chromosomes never complete a second mitosis and the cells succumb to these static states. These experiments reveal that multiple nucleation sites originate that spread to form compacted chromosomes and that the position of the cell cycle when nucleation initiates dictates the final chromosome morphology. These findings argue that the presence or absence of other chromatin factors, such as cohesin, influence PHF13’s ability to drive prophase-like chromosome condensation and that global phase separation of chromatin is occurring.

### PHF13 oligomerizes via its NTD and PHD domains

PHF13 formed filaments that encapsulated the chromosomes resulting in a refractive difference from the nucleoplasm (Figure 2C and E), suggesting PHF13 driven PIPS. To verify PHF13’s oligomerization potential we first performed different *in silico* analysis (Tango, PONDR and AlphaFold) to see if PHF13 contains any intrinsic features capable of self-interaction (Figure 3A–B). Cross analysis of Tango and PONDR, which predict putative aggregation domains (black peaks) and intrinsically disordered regions (IDRs; red peaks) respectively, identified a highly structured aggregation domain at residues 30-40 within the N-terminal domain (NTD) and two weaker aggregation regions bookmarking the PHD domain at residues 232-238 and 272-280 and multiple IDRs distributed throughout PHF13 (Figure 3A). To gain additional structural *in silico* insight into PHF13’s self-interaction potential we employed the recently published AlphaFold2_advanced algorithm, to model PHF13’s ability to homo-dimerize (Figure 3B). Consistent with PONDR, the AlphaFold2 algorithm did not predict structure for most of PHF13, implying that it is predominantly a disordered protein (Figure 3A and B). In addition, however, AlphaFold2 predicted that PHF13 can homo-dimerize via residues 24-40, where Tango predicted a high probability self-aggregation domain (Figure 3A–B). To test these predictions, we performed classical co-immunoprecipitation experiments using differentially tagged PHF13 full length and deletion proteins (Figure 3C). To this end, Flag-PHF13 (1-300aa) or Flag-PHF13 deletion mutants (100-200, 1-150 and 150-300) were co-expressed with EGFP-PHF13 and immunoprecipitated using Flag-M2 agarose (Figure 3C). These experiments revealed that full-length Flag-PHF13, the N-terminal half (Flag-PHF13_1-150) and the C-terminal half (Flag-PHF13_150-300) of PHF13 were all capable of co-precipitating EGFP-PHF13. In contrast, Flag-PHF13_100-200 did not co-precipitate EGFP-PHF13 (Figure 3C), indicating that PHF13 can self-associate via N- and C- terminal interactions, located in the first (1-100) and last (200-300) 100 aa, consistent with an oligomerization potential (Figure 3C). To explore whether PHF13 N- and C- terminal self-interactions are direct or indirect we next performed an *in vivo* FACS-based FRET approach using tagged PHF13 and deletion proteins (Figure 3D). In FACS-based FRET, YFP-PHF13 was co-expressed with various CFP-PHF13 mutant and full-length proteins and evaluated for the excitation of a FRET signal by FACS, indicating a proximity of 10Å or less and implying a direct interaction (Figure 3D). The expression levels and nuclear localization of all fused proteins were controlled by immunofluorescence and immunoblot (Figure S3B-C). In addition, a YFP-CFP fusion protein and co-expression of YFP and CFP and of YFP-HP1α and CFP-HP1α served as positive, negative and biological dimer FRET controls, respectively (Figure 3D). Co-expression of CFP-PHF13 with YFP-PHF13 gave an average FRET signal of ~60% by fluorescent cell sorting, indicating that full length PHF13 can directly self-associate *in vivo* (Figure 3D). Interestingly, this was significantly higher than the FRET signal obtained by co-expression of YFP-HP1α and CFP-HP1α (~30%), consistent with the possibility of PHF13 oligomerization *in vivo* (Figure 3D). In line with *in silico* predictions (Figure 3A and B) and co-immunoprecipitation experiments (Figure 3C), we were able to map a direct interaction to the first 150 N-terminal amino acids and the NTD (21-70 aa; Figure 3D), whereas ΔNTD or Δ24-40 eliminated the FRET signal. In contrast to the N-terminal region, CFP-150-300 did not generate a FRET positive signal with YFP-PHF13 (Figure 3D), arguing that the C-terminal interaction interface in PHF13 is mediated indirectly via other proteins. To refine the mapping of the N- and C-terminal dimerization regions we co-expressed Flag-PHF13 with either YFP-PHF13 N-terminal mutants (Figure 3E) or with YFP-PHF13 C-terminal (Δ7;150-300) mutants (Figure 3F) and looked for their ability to co-precipitate. These experiments confirmed that the NTD and more specifically aa 24-40 mediate PHF13’s N-terminal dimerization (Figure 3E). A weak residual co-immunoprecipitation of Flag-PHF13 was observed for YFP-ΔNTD and YFP-Δ24-40 proteins (Figure 3E) consistent with the existence of a second indirect C-terminal dimerization. Interestingly, deletion of the PHD domain in the C-terminal Δ7 (150-300) protein (Δ7ΔPHD) abrogated the C-terminal dimerization (Figure 3F) as well as cohesin (SMC1 and SMC3) and H3K4me3 interactions (Figure 1D and 3F). In contrast however, point mutations in the PHD domain (Δ7M246A and Δ7W255A) which disrupt PHF13 binding to H3K4 methylated histones did not impair its C-terminal dimerization with Flag-PHF13 nor its cohesin interaction (Figure 1D and 3F). These findings demonstrate that the PHD domain mediates the indirect C-terminal dimerization and implicate cohesin as a connector protein (Figure 3F). To further validate PHF13’s oligomerization potential and to approximate the size of PHF13 homo-oligomers *in vitro*, we performed size exclusion column chromatography (Superose 6 10/300) of purified recombinant PHF13 and deletion proteins after the proteolytic removal of the GST tag (Figure 3G and S3C). Based on calibration standards, monomeric PHF13 (MW=34kDa) was expected to elute around fraction 36 on a Superose 6 column. However, PHF13 and PHF13-ΔPHD eluted in fractions 22 to 29, with the most predominant peak detected in fraction 25 and only a minor monomeric peak at fraction 36 (Figure 3G and S3C). This indicates that recombinant PHF13 and PHF13-ΔPHD preferentially exists as oligomers (between ~400-700 kD in size) and that the PHD domain does not mediate direct PHF13-PHF13 interactions. In contrast, deletion of the NTD or residues 24-40 reduced recombinant PHF13 to predominantly monomeric and dimeric fractions (34 to 38), indicating that these 16 residues mediate direct self-interactions (Figure 3G and S3C). Similar results were also obtained using *in vitro* translated PHF13 and PHF13ΔNTD proteins on a Superdex 200 column (Figure S3D). Together, these results reveal that PHF13’s conserved NTD is a homo-dimerization domain and demonstrate PHF13’s ability to oligomerize via direct N-terminal and indirect C-terminal interactions, consistent with the expectations of PIPS.

**Figure 3.**
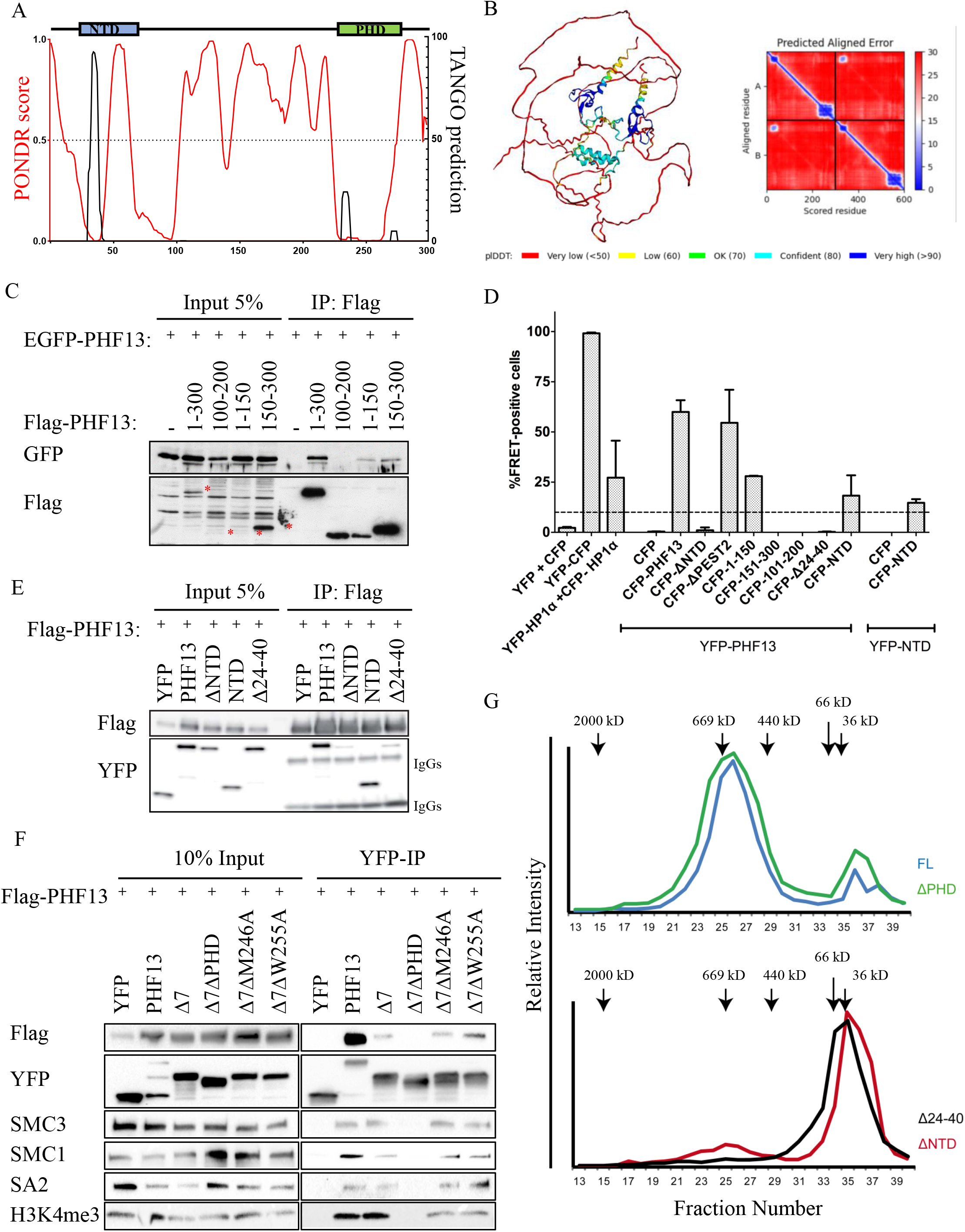
PHF13 oligomerizes via its NTD and PHD domains. (A) A merged PONDR (Red) and Tango (black) in silico analysis of PHF13. (B) AlphaFold2_advanced analysis of PHF13 dimers (shown as a string model and contact map). The confidence of the called structure is colour coded. (C) Co-IP of Flag-PHF13 and Flag-PHF13 deletion proteins with EGFP-PHF13. FACS-FRET analysis of 293 cells co-transfected with YFP and CFP fusion proteins. 10% FRET signal was arbitrarily defined as the minimum threshold and is denoted by a dotted line. (E) Flag IP: WB from lysates co-expressing Flag-PHF13 with YFP, YFP-PHF13 and YFP-PHF13 deletion proteins. (F) GFP-Trap:WB from cells co-expressing Flag-PHF13 with YFP, YFP-PHF13 or C-terminal YFP-PHF13 mutant proteins. WB: were probed for GFP and Flag (C, E and F) and for the co-precipitation of the endogenous cohesin complex and H3K4me3 (F). (G) Graphical representation of the size exclusion elution profile (Superose 6 10/300) from GST-cleaved recombinant PHF13 full-length (FL; blue) and deletion proteins (ΔPHD; green, ΔNTD; red and Δ24-40; black), obtained by dot blotting of individual fractions and quantification by Image quant.

### The N- and C-terminal dimerization domains of PHF13 are essential for condensation

To validate the role of PIPS in PHF13 induced chromatin compaction, we tested the impact of N- and C- terminal dimerizing mutant proteins expressed at similar levels on PHF13’s ability to condense chromatin and on its chromatin avidity (Figure 4A–J and S4A and B). Nucleation of PHF13 induced condensation originated from nucleoli (Figure 2F and S2E) and coincides with the mitotic phenomenon (Leung et al., 2004) of nucleolar dissociation (Figure 4A and S4C), allowing us to use nucleolar integrity (NOH61, Ki67, Nucleolin and Topoisomerase I) as a second proxy for condensation. To this end, we found that deletion of PHF13’s N-terminal domain (ΔNTD) or its PHD domain (ΔPHD) abolished its ability to compact chromatin (Figure 4B and C) indicating that their specific molecular functions are required for this phenotype. Consistent with N-terminal dimerization being the important feature, Δ24-40 also failed to compact chromatin (Figure 4D) whereas deletion of the PEST1 (ΔPEST1) domain (aa 50-88) which partially overlaps with the NTD (aa 21-70; Fig 1A and S1A) but retains the homo-dimerization motif (aa 24-40), still compacted chromatin (Figure 4E). Furthermore, we could rescue the condensation defect of the PHF13ΔNTD protein by fusing it to the exogenous dimerization motif of GAL4 ^DD^ (50-147aa), confirming the importance of PHF13 oligomerization in driving higher chromatin order (Figure 4F). However, the question remained whether the PHD domain is important for this phenotype due to its ability to tether PHF13 to histones or due to its role in C-terminal dimerization via cohesin and possibly other substrates. To address this, we expressed PHF13 PHD domain point mutants (M246A or W255A) which disrupt H3K4me3 binding but still dimerize and interact with cohesin (Figure 1D) to see whether these single point mutations were sufficient to inhibit PHF13’s ability to compact chromatin (Figure 4G and H). In contrast to ΔPHD, PHF13M246A and PHF13W255A still compacted chromatin as can be seen in the differential interference contrast (DIC) images and by the disruption of the nucleoli marker NOH61 (Figure 4G and H). Nevertheless, the condensation was fundamentally different for the PHF13 PHD point mutant proteins, as they resulted in compacted chromatin aggregates and not chromosome like structures, arguing that PHF13’s association with H3K4me3 is important for correct compaction morphology but not condensation per se. All together, these findings support that the oligomerization mediated by PHF13’s N- and C- terminal domains are essential features for PHF13 induced condensation.

**Figure 4.**
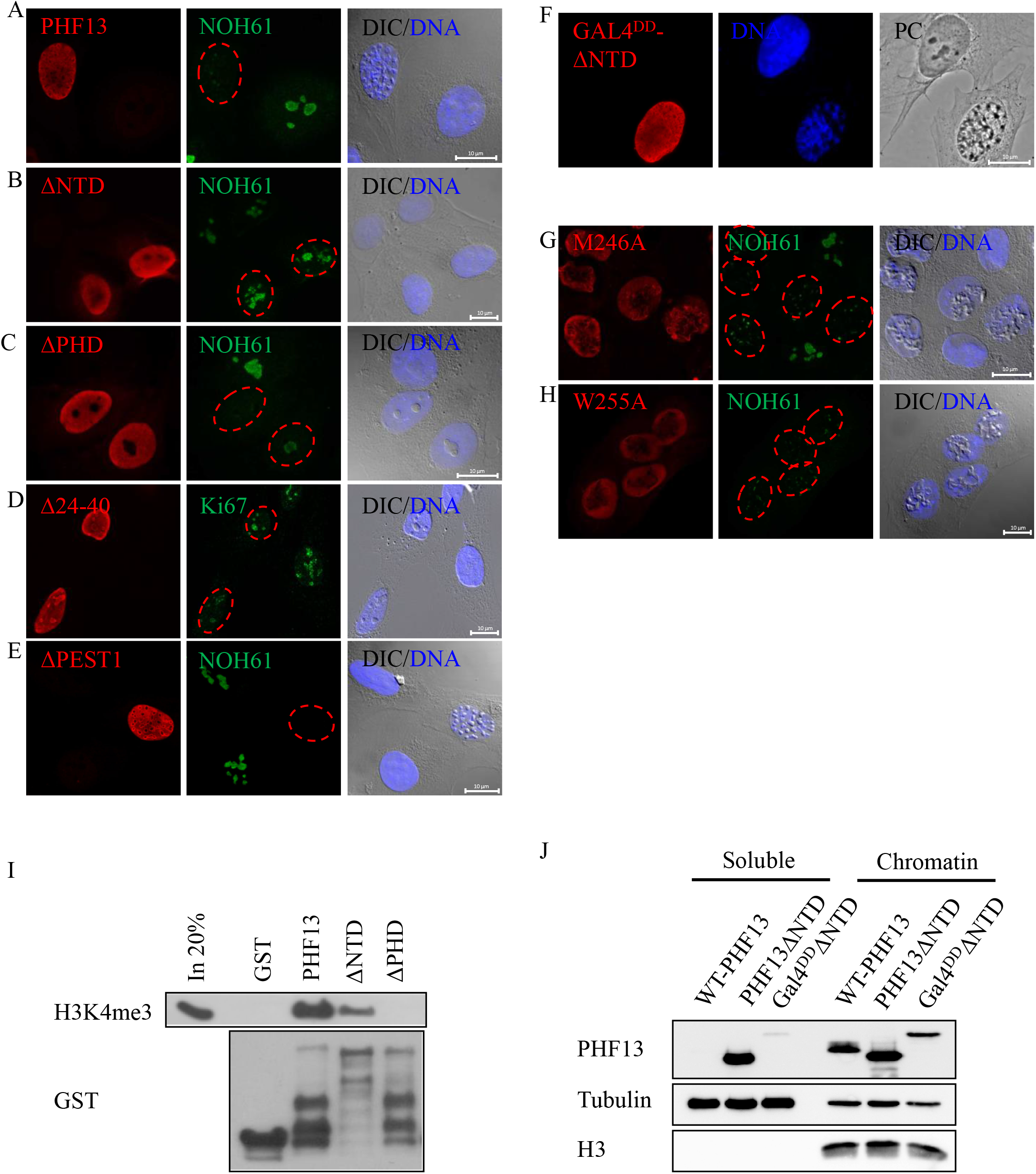
PHF13 oligomerization is necessary for its ability to compact chromatin. (A-C) IF staining of U2OS cells transfected with 1μg of PHF13, PHF13ΔPHD, PHF13-M246A and PHF13-W225A (A), PHF13ΔNTD, PHF13Δ24-40 and PHF13ΔPEST1 (C), and PHF13-GAL4^DD^ΔNTD (C). PHF13 was stained with rat mAb 6F6 (A) or 1D3 (B and C) and DNA was stained with Hoechst. Nucleoli (green) were detected with NOH61 or Ki67 antibodies (A and B). Differential interference contrast (DIC) and phase contrast (PC) images were captured on a Zeiss Confocor 2 confocal microscope (A and B) and on a Zeiss Axiophot Phase Fluorescence microscope (C). (D) GST-pulldowns of nuclease digested chromatin lysates immunoblotted for GST and H3K4me3. (E). WB of fractionated lysates from cells expressing PHF13, PHF13ΔNTD and PHF13-GAL4^DD^ΔNTD and detected with PHF13 (rat mAb 6F6), Tubulin and H3 antibodies.

The ability of PHF13 to oligomerize via direct N-terminal and indirect C-terminal interactions creates polyvalent PHF13 chromatin interactions, with alternating DNA and histone binding domains. This extended chromatin valence is predicted to strengthen PHF13’s chromatin binding avidity and facilitate its ability to span/bridge multiple nucleosomes. To test this, we looked for the ability of equimolar amounts of recombinant PHF13, PHF13ΔNTD and PHF13ΔPHD to precipitate H3K4me3 from chromatin nuclear lysates (Figure 4I) and for the chromatin localization of oligomerization competent (PHF13 and GAL4^DD^-PHF13ΔNTD) and incompetent (PHF13ΔNTD) proteins (Figure 4J). GST-PHF13 efficiently precipitated H3K4me3 an interaction that is lost in the GST-PHF13ΔPHD pulldown and that is absent in the GST-control (Figure 4I) as previously reported (Chung et al., 2016), whereas deletion of PHF13’s NTD, resulted in a reduced precipitation of H3K4me3 (Figure 4I) in line with PHF13 oligomerization increasing its chromatin avidity. Consistently, immunoblotting of fractionated lysates from wild-type PHF13, PHF13ΔNTD and GAL4^DD^-PHF13ΔNTD demonstrated that deletion of PHF13’s NTD caused a substantial shift of PHF13 from the chromatin fraction to the nucleoplasmic fraction, which could be recovered by fusion of PHF13ΔNTD to the Gal4 dimerization motif (Figure 4J). Together, these data demonstrate that PHF13’s ability to oligomerize, increases its chromatin valence and avidity, and is essential to promote higher chromatin order.

### PHF13 and cohesin depletion impair condensation and cause mitotic cell death

Since PHF13 interacts with cohesin via its PHD and both promote higher chromatin order changes, we reasoned that they might cooperate in this function at mitotic transition. To provide functional evidence for this possibility, we synchronized cells depleted of PHF13 and/or SMC3 at the G2/M border and released them into mitosis (Figure 5A–E). We then followed their release using time-lapse microscopy, live cell SPY-DNA stain and bright field imaging (Figure 5A and S5A) and quantified the mitotic index, chromatin compaction and mitotic death in single cells across time points and conditions (Figure 5B–D and S5B). We observed that PHF13 and SMC3 depletion alone or in combination greatly reduced the mitotic index (Figure 5A; black circles and B) and impaired condensation (Figure 5C and S5B). Furthermore, co-depletion of PHF13 and SMC3 led to a significant increase in mitotic cell death (Figure 5D). Immunoblotting analysis confirmed the efficient depletion of PHF13 and SMC3 which corresponded with lower H3S10 phosphorylation levels (Figure 5E) reflecting the decreased mitotic index (Figure 5B). Furthermore, cyclin B1 was significantly reduced in PHF13/SMC3 co-depleted cells, further implicating an impaired mitotic entry (Figure 5E). To get a clearer picture of the impact on mitotic condensation, we probed synchronized and fixed cells for H3S10 phosphorylation and captured images at high magnification (Figure 5F). Consistent with live cell analysis, fewer cells were labelled with H3S10p in PHF13si and/or SMC3si conditions in comparison to controls (Figure S5C) consistent with a reduced mitotic index. Interestingly however, high magnification of H3S10p stained cells in the PHF13si and SMC3si only conditions, failed to reveal a notable impact on mitotic chromatin compaction whereas PHF13 and SMC3 co-depleted cells lacked chromosome definition and looked apoptotic (Figure 5F). Together, these findings are consistent with PHF13 and cohesin having complementary functions in mitotic higher chromatin order and mitosis.

**Figure 5.**
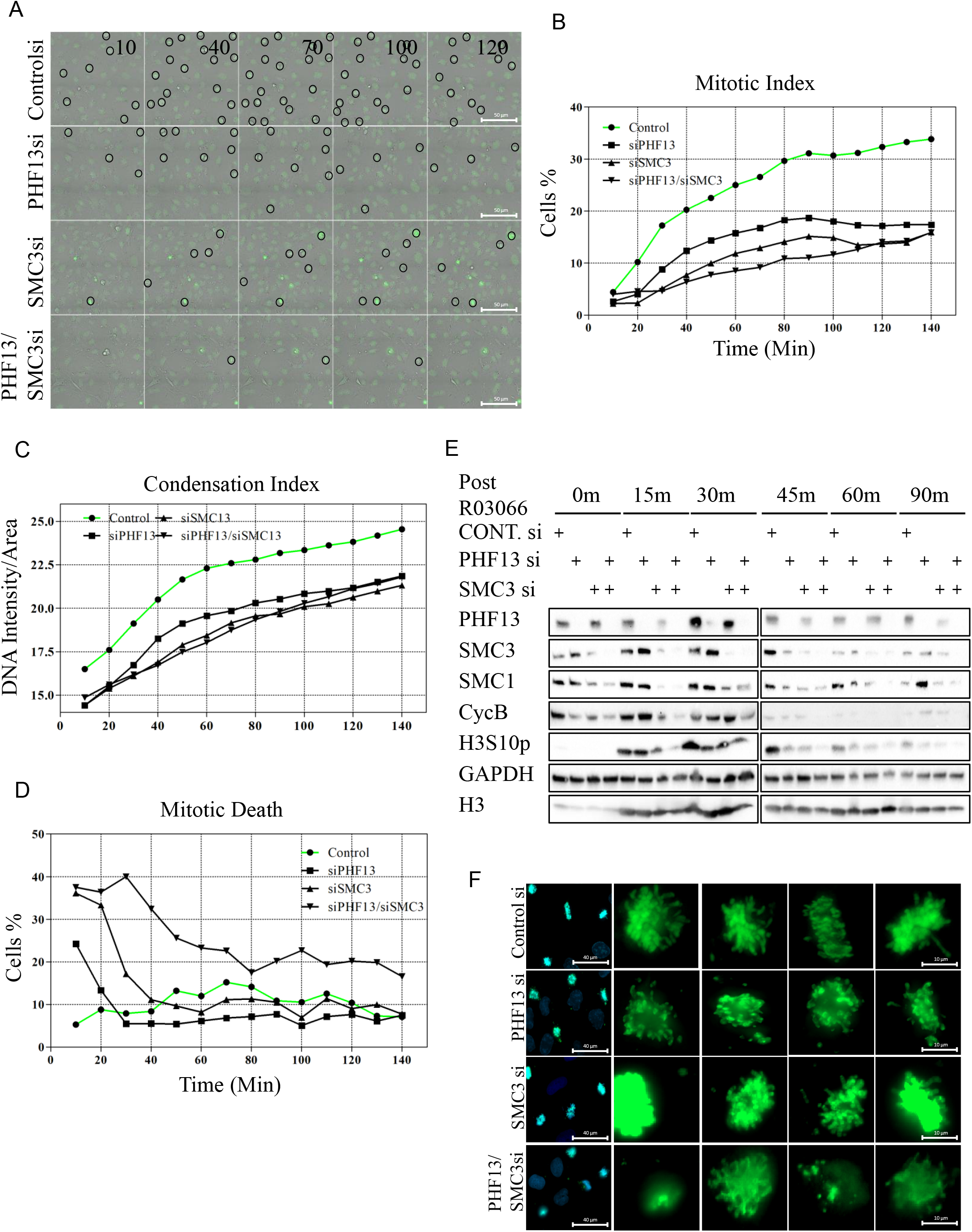
PHF13 and SMC3 depletion impair mitotic condensation and cell division. (A) Live cell microscopy of synchronized wild-type (cont. siRNA), PHF13 siRNA, SMC3 siRNA and PHF13/SMC3 siRNA depleted conditions. Mitotic cell (black circles), dead cells (red circles). Bright field and DNA images were captured on a Zeiss cell discover. (B-D) The mitotic index (B), chromatin compaction (C) and mitotic cell death (D) was determined using machine learning and Zeiss software. Values were plotted over time and is presented as the averages from two biological and two technical replicates. (E) WB of synchronized lysates probed with specific antibodies to SMC3, SMC1, PHF13, cyclin B, GAPDH, H3S10P and H3. (F) High magnification light microscopy images of fixed G2/M synchronized and released cells (75m) stained with H3S10p to evaluate chromosome morphology. Images were taken with a Zeiss observer at 20x and 100x magnification.

### Molecular Dynamics Simulations recapitulate our *in vivo* observations

To computationally investigate the folding mechanism driving PHF13 mediated chromatin compaction, we employed a polymer physics model based on the Strings and Binders Switch (SBS) model (Barbieri et al., 2012; Chiariello et al., 2016) and the Transcription Factors (TF) model (Brackley et al., 2013). In this model, a chromatin filament is represented by a chain composed by *N* beads, equipped with specific binding sites that can interact with molecules, which populate the surrounding environment (Figure S6A and B). In our simulations, we use polymers with *N* = 2576. By imposing a size of 25kb for each bead, the resulting length equals the size of human chromosome 20 (~ 64.4 Mb). The model for the PHF13 protein consists of a small molecule made of three distinct domains, corresponding to the PHD domain, NLS domain and the N-terminal domain (Figure 6A). PHF13 proteins are at a concentration *c*, expressed as volume fraction. PHF13 interact with chromatin through a strong attractive interaction between the PHD domain and H3K4me3 sites (*E*_*PhD,H*3*K*4*me*3_) and weaker attractive interaction between PHD domain with H3K4me1/2 sites (*E*_*PhD,H*3*K*4*me*1/2_). Binding sites are regularly located along the polymer chain with density of ~0.1 (1 site every 11 beads) for H3K3me3 and ~0.4 for H3K4me1/2, according to the scheme shown in Figure S6A. A non-specific, weak interaction is also present between NLS and all the polymer beads (*E*_*NTD,NTD*_). Finally, PHF13 molecules can attractively interact with themselves through an NTD-NTD (Figure 6A) interaction (*E*_*NLS,polymer*_). In addition, we model the binding of cohesin dimers with the polymer and restrict its binding to H3K4me3 and CTCF sites (located with a density of ~0.1 and positioned beside H3K4me3), consistent with genome-wide profiles (Figure 1B). Finally we model PHF13 with cohesin (E_PHD, Cohesin_), allowing PHD domain-cohesin interactions, based on the current findings of this paper (Figure 1, 3 and S6B). Overall, the described system includes both protein-chromatin and protein-protein interactions (Figure 6A), and the resulting equilibrium structure depends on the interplay between these interactions (Chiariello et al., 2020). The Molecular Dynamics (MD) simulations demonstrate that PHF13 is able to phase separate chromatin in the absence of cohesin, resulting in a collapse globule at the end state equilibrium (Figure 6C–D and S6C). Furthermore, by shutting off either the NTD or PHD domain interactions (Figure 6C and D), PHF13 loses its ability to phase separate the chromatin fiber, nicely recapitulating our *in vivo* findings and supporting that PHF13 linearly compacts chromatin via its ability to interact with itself and form multivalent chromatin interactions. Modelling PHF13 and cohesin together, resulted in compacted rod-like chromosome structures at end-state equilibrium (Figure 6F–G and S6C), whereas modelling cohesin only, induced chromatin loops and resulted in a more elongated phase separated polymer (Figure 6E and S6D). Interestingly, when all interactions were allowed except for PHF13’s PHD interactions with H3K4 methylated histones, phase separated aggregates were still observed. However they formed random configurations, similar to what was seen *in vivo* (Figure 4G–H and S7A) indicating that PHF13 binding to H3K4 methylation is important for determining the overall shape of chromatin fiber but not for its phase separation propensity. These simulations recapitulate our *in vivo* findings and support a two-step polymer-polymer phase separation process where linear compaction driven by PHF13 and bridging induced compaction mediated by cohesin cooperate to produce rod-like chromosomes.

**Figure 6.**
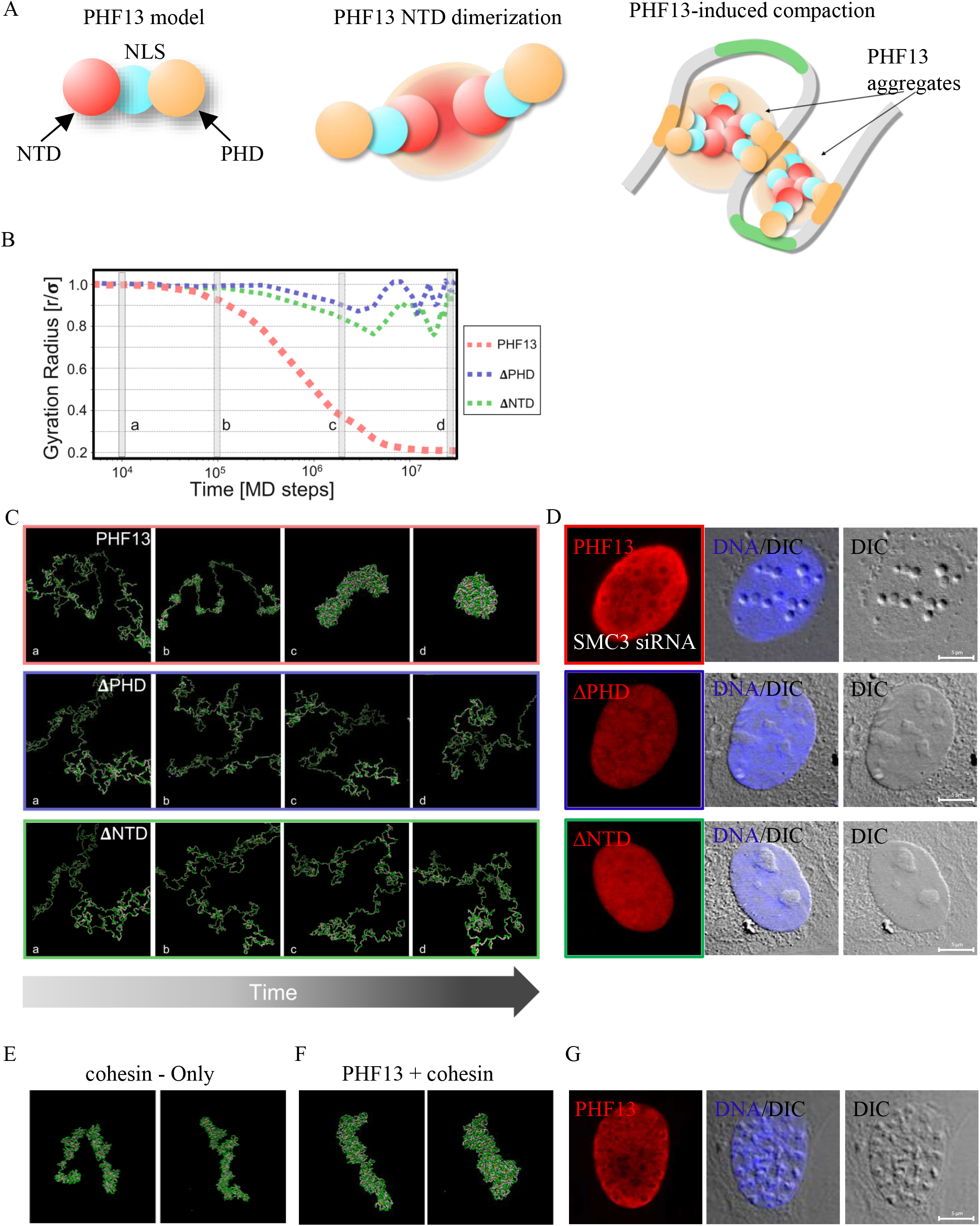
Molecular dynamic simulations. A) We model PHF13 as a molecule made of 3 beads representing the NTD, NLS and PHD. NTD mediates PHF13 self-interaction and the PHD domain mediates an interaction with chromatin and with cohesin. All interactions cooperatively contribute to chromosome condensation. B and C) Polymer folding dynamics from initially randomly open configurations is monitored by the gyration radius. When all PHF13 interactions are enabled and cohesin is not considered, the polymer stably folds in to compacted structures and the gyration radius sharply decreases (PHF13, red curve). Deletion of PHD (ΔPHD, blue curve) or deletion of NTD domain (ΔNTD, green curve) prevent polymer compaction, which remains in an open conformation. E and F) Polymer model end states of cohesin-only or PHF13+cohesin. D and G) IF images of U2OS cells over expressing PHF13, ΔPHD and ΔNTD in the absence (SMC3 siRNA-where noted) or presence of endogenous cohesin.

## Discussion

In this study, we show that PHF13 and cohesin cooperate in higher order chromatin transition and we propose that polymer-polymer phase separation (PPPS) is the underlying mechanism of this process. While the role of liquid-liquid phase separation in interphase chromatin architecture and function is widely referenced with a multitude of examples (Hnisz et al., 2017; Kilic et al., 2019; Strom et al., 2017), to date there are no *in vivo* biological examples of PPPS to the best of our knowledge, nor a clear understanding of the mechanism of mitotic condensation. Our findings provide compelling *in vivo* support that chromosome condensation follows the biophysical principles of PPPS. In contrast to LLPS, which is driven typically by weak multivalent IDR based protein-protein interactions formed in the presence or absence of chromatin, PHF13 forms ordered self-interactions via its aggregation domains, apparently independent of its IDRs, resulting in a polyvalent chromatin effector protein with increased chromatin avidity, supporting that PHF13 induced condensation involves strong intermolecular protein-protein and protein-chromatin contacts. Notably, the compacted chromosomes display a refractive difference from the nucleoplasm and are encapsulated by a PHF13 polymer. These features are in line with polymer-polymer phase separation models, which require stable multivalent protein nucleic acid interactions (Erdel and Rippe, 2018). Based on our findings we propose the following mechanism and model (Figure 7) to explain chromatin to chromosome transitions. In general, PHF13 oligomerizes facilitating multivalent chromatin association and promoting a linear compaction of chromatin. In addition, PHF13 interacts with cohesin, which drives bridging induced compaction, synchronizing these chromatin forces and satisfying two-step condensation models (Figure 7).

**Figure 7.**
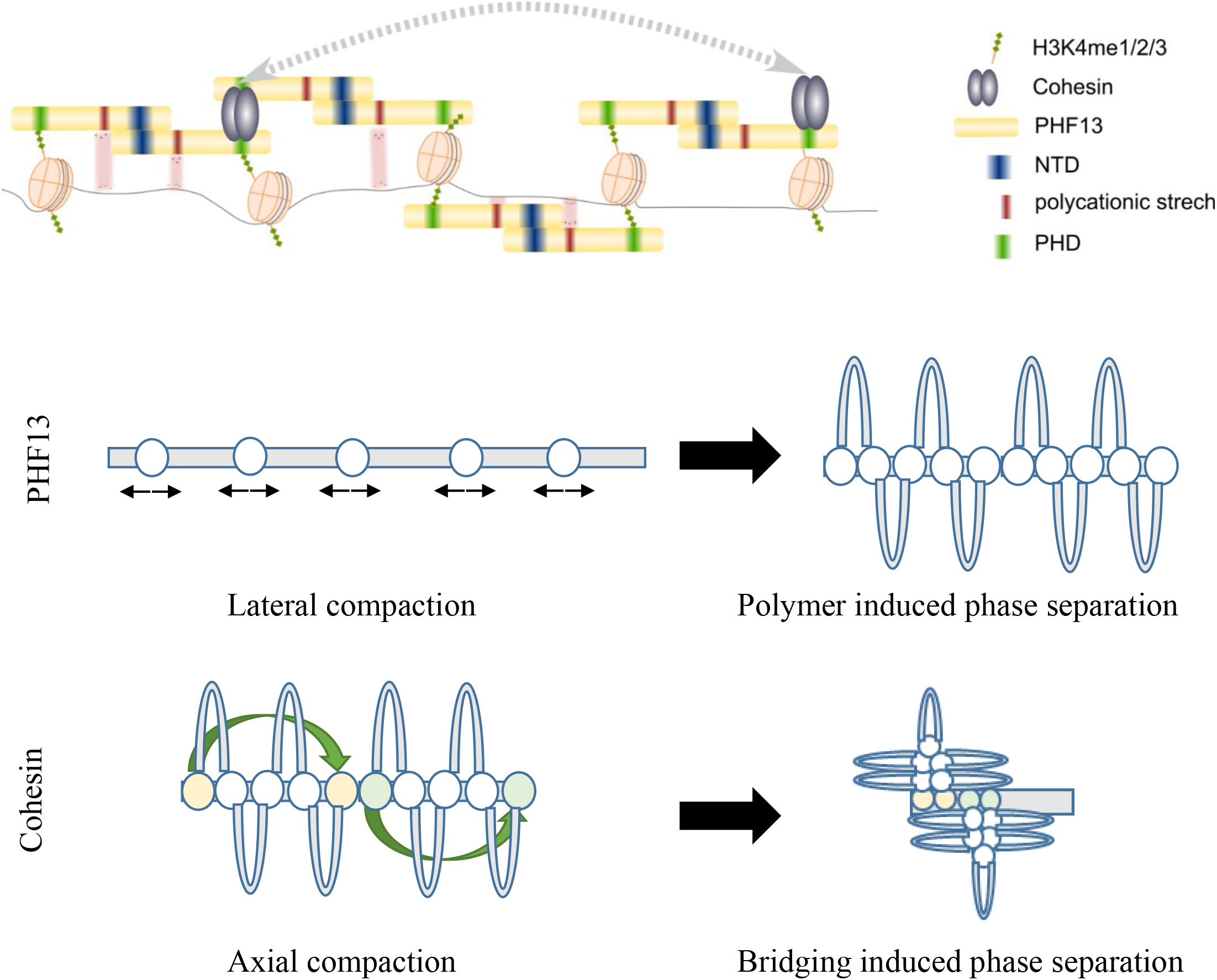
Model of PHF13 and Cohesin induced condensation. In the upper part of the model, PHF13 (denoted as a yellow bar) is able to oligomerize via direct N-terminal interactions (blue) and indirect C-terminal interactions (green) with cohesin (grey spheres). This oligomerization creates a PHF13 polymer with alternating DNA binding (red; cation π interactions) and H3K4me3 binding (green; PHD domain) regions allowing it to spread along chromatin fiber and facilitate linear compaction via polymer induced phase separation. Cohesin forms longer distance loops and can induce bridging induced phase separation (lower part of image).

In support of their cooperative role and a two-step condensation process, Molecular Dynamics simulations modelling PHF13 and cohesin’s ability to compact a chromatin polymer resulted in rod-like end-states when both were acting on the chromatin polymer and when they interacted with each other. Shutting off the interaction of PHF13 with cohesin, via silencing the PHD domain, resulted in a more open elongated chromatin structure similar to cohesin-only (Figure 6E and S7), whereas silencing cohesin and allowing only PHF13 resulted in a collapse globule (Figure 6C and S6C) which could be recapitulated in vivo by expressing PHF13 in SMC3 depleted cells (Figure 6D). Consistently, only cells expressing a critical concentration of PHF13 in S-phase or later were able to condense chromatin into chromosomes, whereas cells expressing a critical concentration of PHF13 in G1-formed chromatin aggregates. These observations suggest that spatiotemporal events such as the establishment of chromosome territories and the cohesion of sister chromatids are required for prophase-like chromosome condensation. In the absence of these events, PHF13 glues chromatin together in a linear and inter-chromosomal fashion forming aggregates. Further speaking in favor of their cooperatively, only the co-depletion of PHF13 and SMC3 was sufficient to abrogate chromosome condensation and trigger mitotic catastrophe. We suggest that in vertebrates, PHF13 and cohesin establish inter-chromatid compaction along the axis and periphery of sister chromatids to promote early prophase condensation with non-resolved chromatids.

We suspect that PHF13 has a redundant role in linear compaction of chromatin based on the fact that PHF13 knockout cell lines exist. It will be interesting to see which other proteins have a homologous function and can compensate for PHF13 and to see what regulates the dynamics and equilibrium of PPPS driven chromatin condensation. It seems reasonable that phosphorylation gradients may influence which factors bind to chromatin and in which constellation to fine-tune PPPS during mitosis. Interestingly, it was recently shown that H3T3 phosphorylation, a modification occurring in late G2 along chromosome arms (Dai et al., 2005), increases PHF13’s PHD binding to H3K4me3 (Jain et al., 2020), suggesting a potential explanation for PHF13’s increased chromatin association at mitotic transition. Finally, it will be important to evaluate in the future the interphase relationship of PHF13 with cohesin and the role of PHF13 in cohesin chromatin recruitment.

## Acknowledgments

We would like to thank Michael Schindler, Carina Banning and Kristin Höhne for their technical support with the FACS-FRET assay, Marcos Torroba for his help with formatting the videos, M. Schmidt-Zachmann for the NOH61 antibody and Theres Schaub and Kirsten Reumann for their help with characterizing PHF13 antibodies and generating plasmids. This work was supported by the VW-Stiftung #97131 and the Max Planck Gesellschaft.

## Author Contributions

SK, FR, RB, TS, HS, AG and AMC performed the experiments. LG performed the informatics. SK, HW, and MV were involved in conceptualizing and managing the project. SK wrote the manuscript.

