## supplementary Figures for "Connecting the Dots: PHF13 and cohesin promote polymer-polymer phase separation of chromatin into chromosomes"

Supplementary Figure 1:

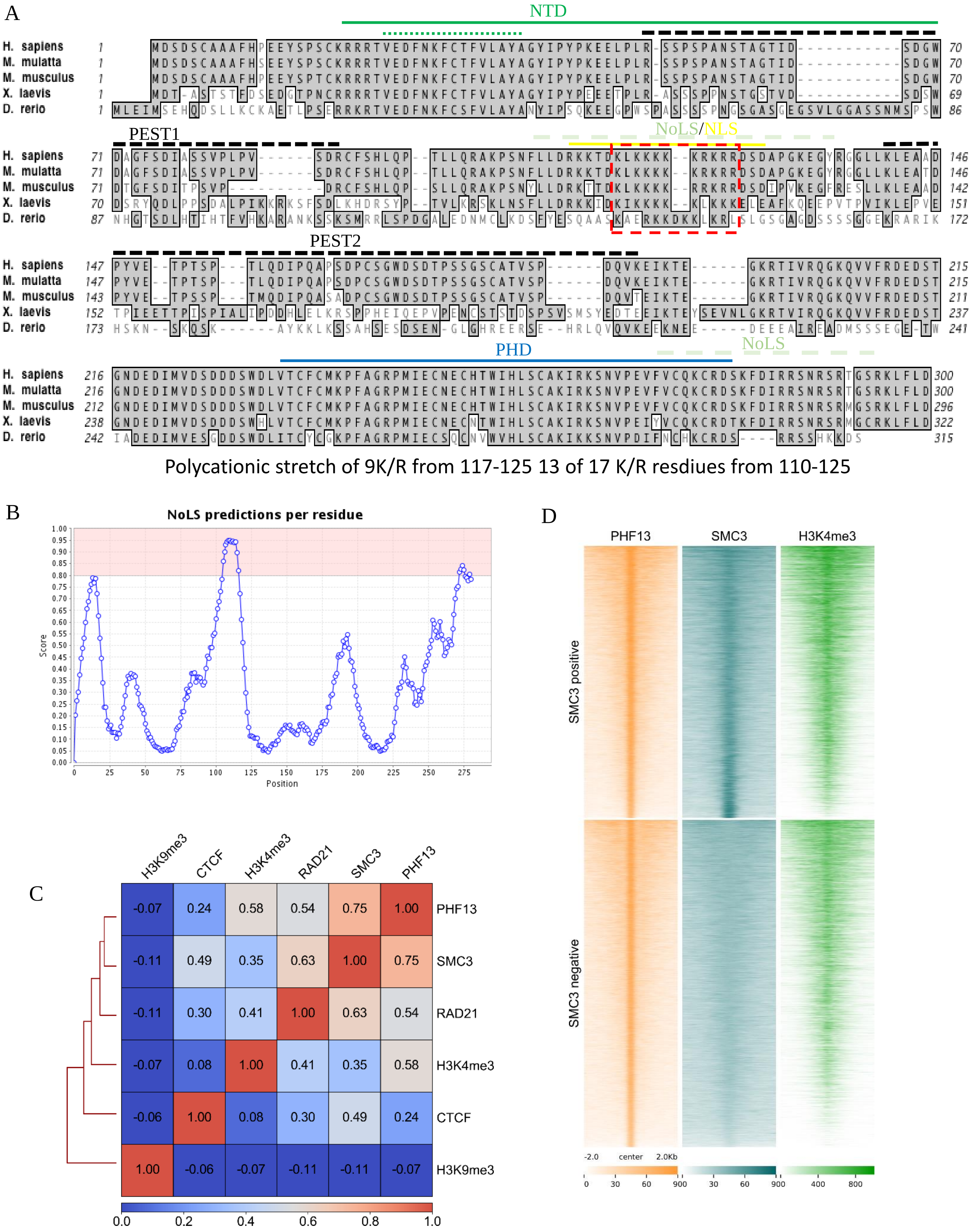

Supplementary Figure 2:

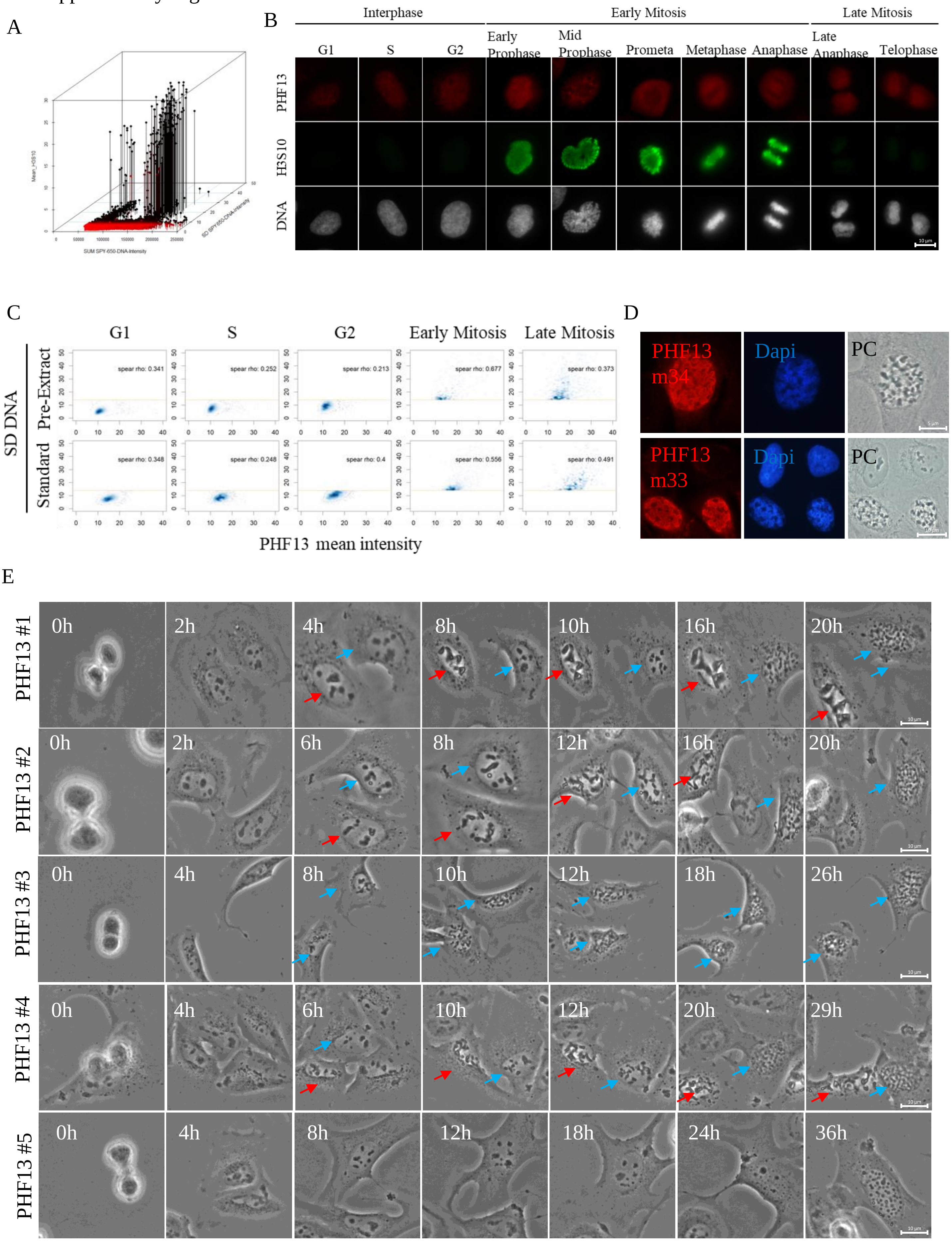

Supplementary Figure 3:

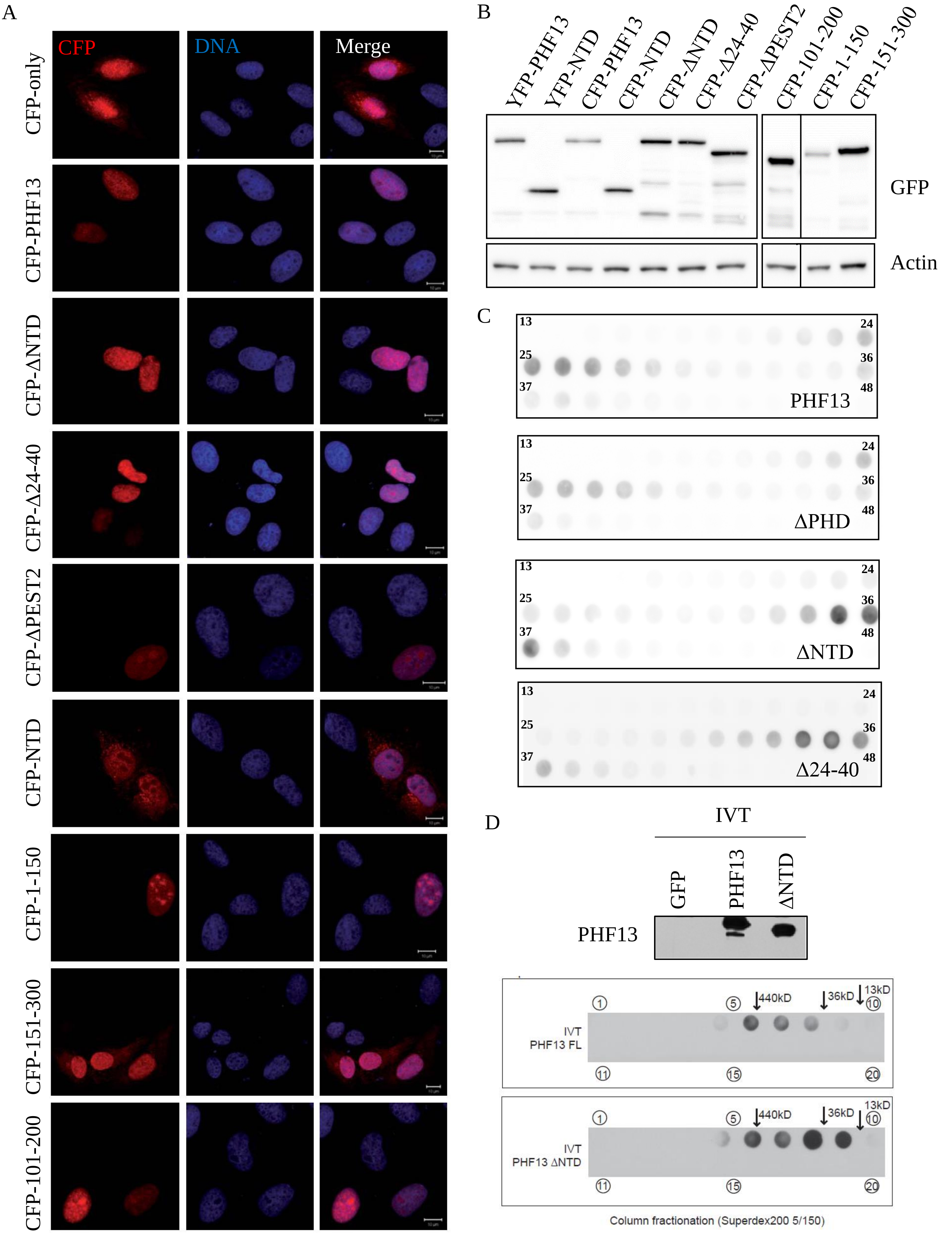

Supplementary Figure 4:

A

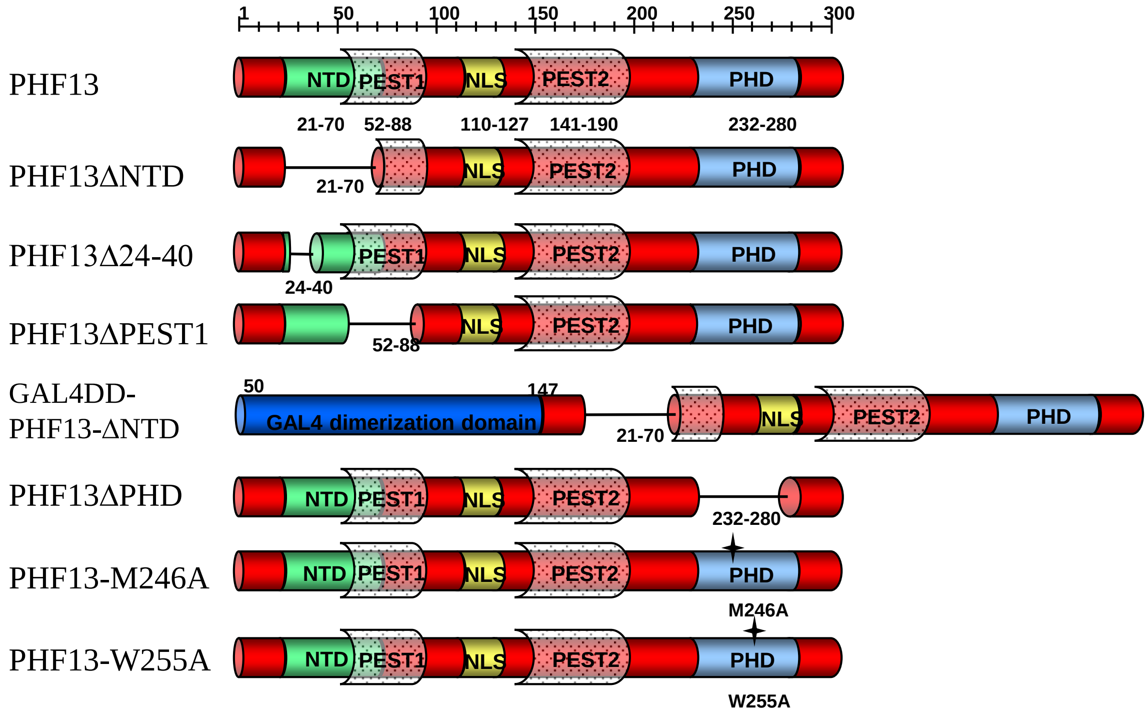


B

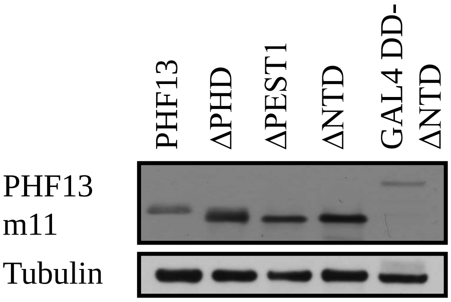


C

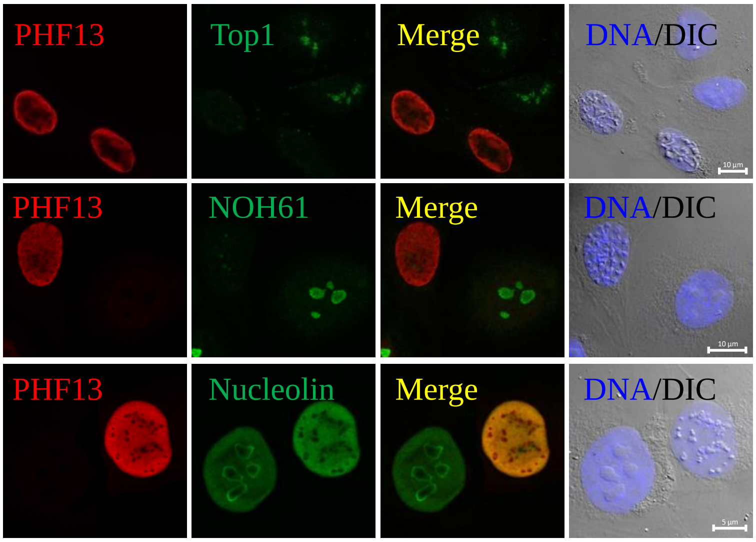

Supplementary Figure 5:

A

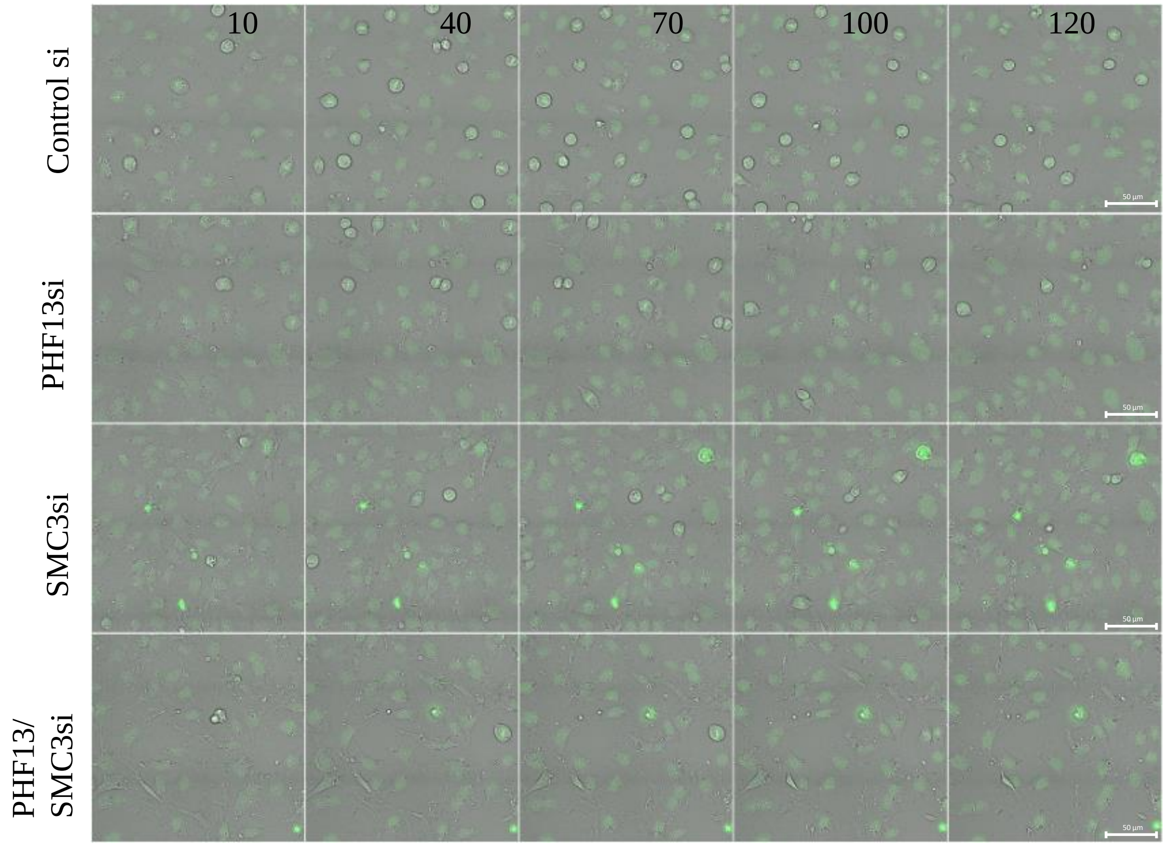


B

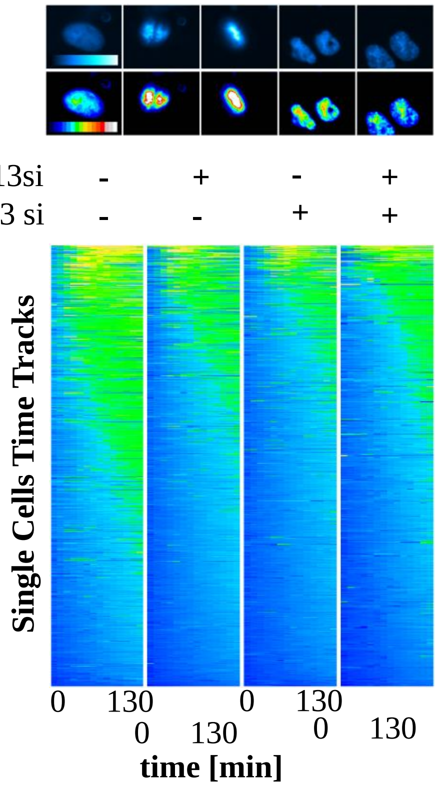


C

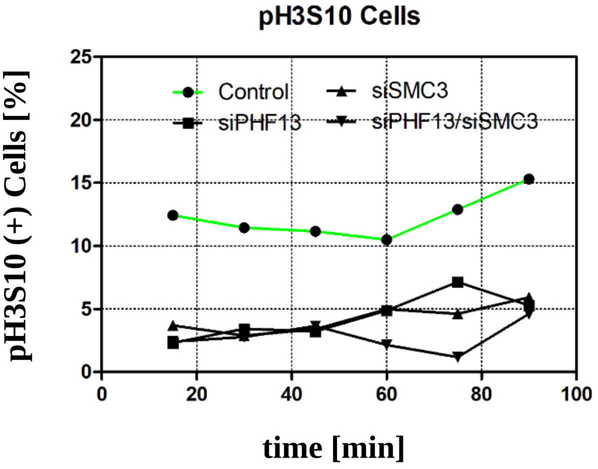

Supplementary Figure 6:

A

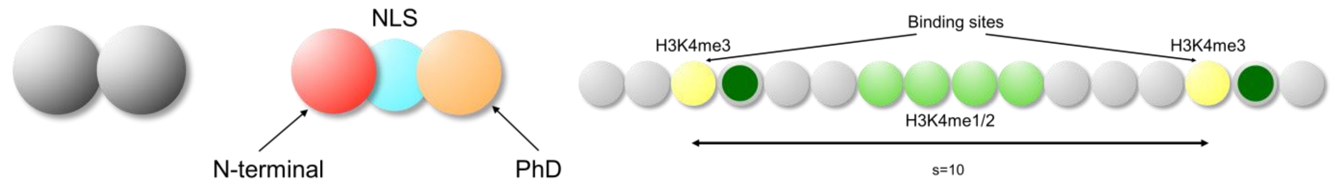


B

Interactions

| Type | | Binding affinity [KbT units] |
|---|---|---|
| NTD-NTD | | 4.1 |
| NLS-chromatin | | 2.4 |
| PHD-H3K4me3 | | 6.1 |
| PHD-H3K4me1/2 | | 2.9 |
| PHD-Cohesin | | 4.1 |
| Cohesin-CTCF | | 8.1 |

C

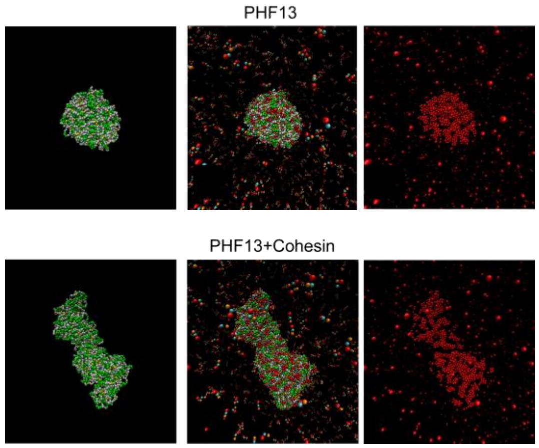


D

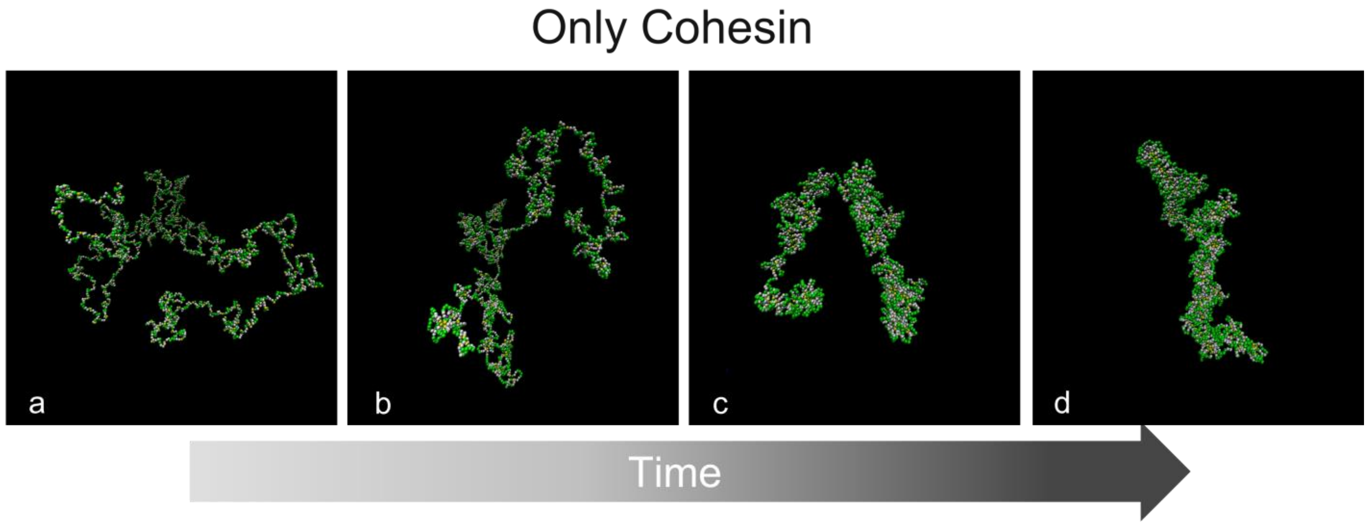

Supplementary Figure 7:

A

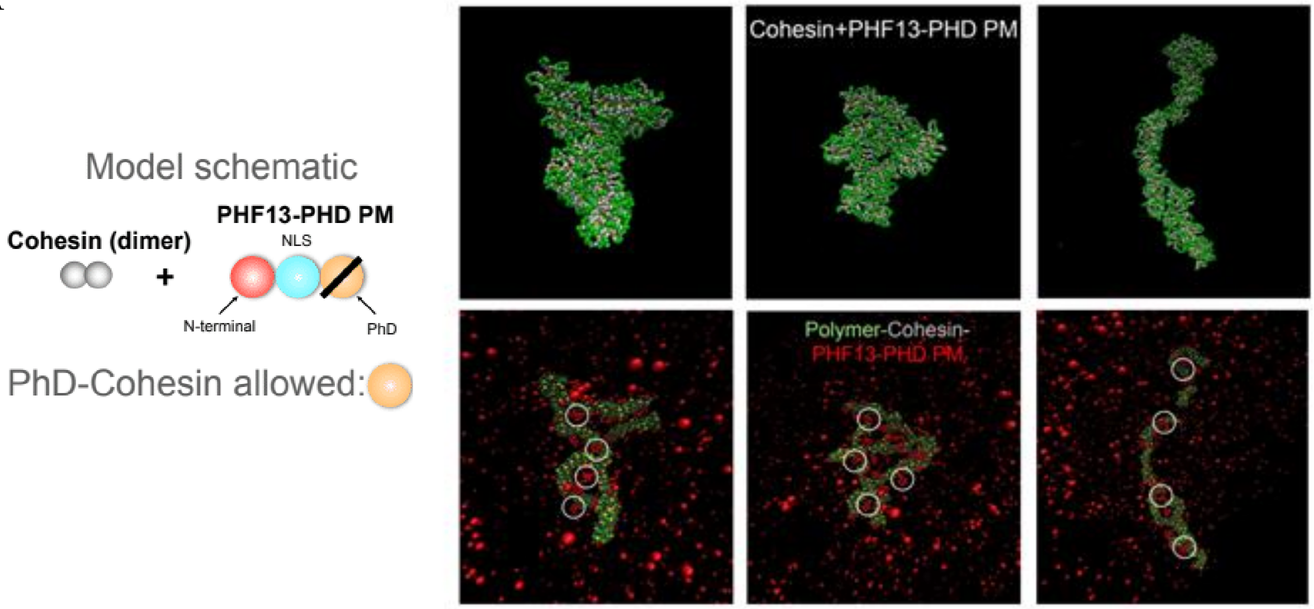


B

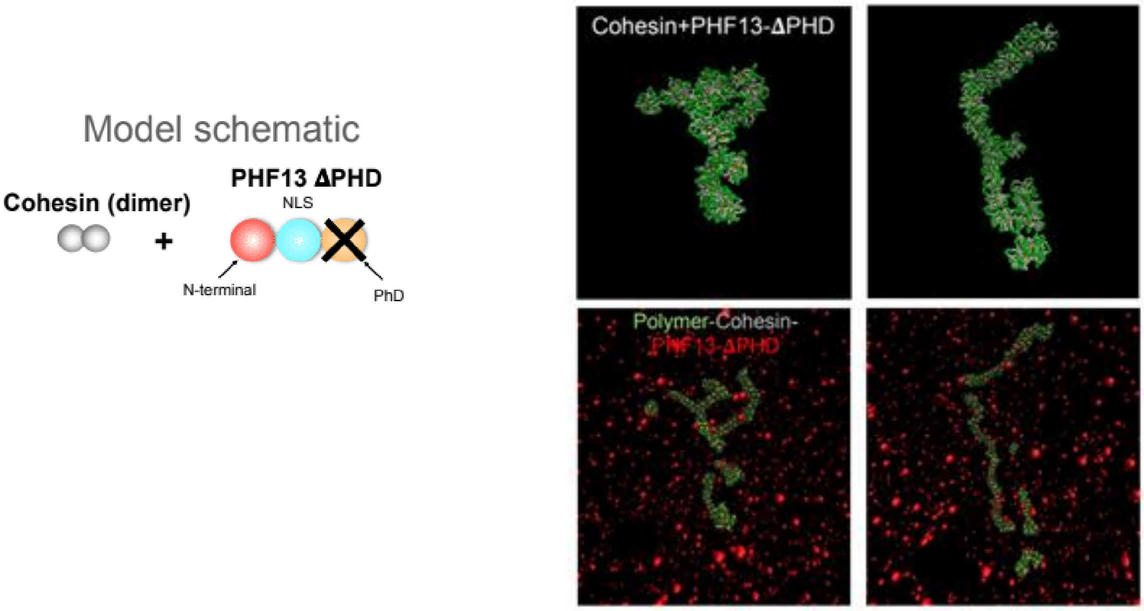
