## Supplementary material for "Connecting the Dots: PHF13 and cohesin promote polymer-polymer phase separation of chromatin into chromosomes": Materials and Methods

**Tables with Titles:**
**STAR Methods:**
**RESOURCE AVAILABILITY**
***Lead Contact***

**Materials availability**
Plasmids generated in this study will be deposited to addgene.
pEGFP-C1-PHF13
pFlag-CMV4-PHF13
pFlag-CMV4-PHF13(100-200)
pFlag-CMV4-PHF13 (1-150)
pFlag-CMV4-PHF13 (150-300)
pEYFP-C1-PHF13
pEYFP-C1-PHF13_NTD(21-70)
pEYFP-C1-PHF13ΔNTD(del21-70)
pEYFP-C1-PHF13del24-40
pEYFP-C1-PHF13(150-300)
pECFP-C1-PHF13
pECFP-C1-PHF13ΔNTD (del21-70)
pECFP-C1-PHF13ΔPEST2 (del141-190)
pECFP-C1-PHF13(1-150)
pECFP-C1-PHF13(150-300)
pECFP-C1-PHF13(100-200)
pECFP-C1-PHF13del24-40
pECFP-C1-PHF13_NTD (21-70)
pGEX-4T3-PHF13
pGEX-4T3-PHF13ΔNTD(del21-70)
pGEX-4T3-PHF13ΔPHD
pCDNA-4TO-PHF13
pCDNA-4TO-PHF13ΔNTD (del21-70)
pCDNA-4TO-PHF13del24-40
pCDNA-4TO-PHF13ΔPEST1(del52-88)
pCDNA-4TO-PHF13M246A
pCDNA-4TO-PHF13W255A
pCDNA-4TO-GAL4$^{DD}$PHF13ΔNTD

**Data and code availability**
A final customer R script row bound 500 individual tracks per condition and plotted the data in matrices using Topo. Colors as look up tables and stored data tables. All here stated scripts/snippets are public available https://github.com/ReneBuschow/SK_2021

**EXPERIMENTAL MODEL AND SUBJECT DETAILS**
Mammalian cell lines: The cell lines used in this study were not authenticated. All cell lines tested negative for mycoplasmas.
***U2OS*:** HTB-96; Human epithelial Osteo-Sarcoma cells (female); wild type p53 and Rb; p16-negative: were grown under standard culture conditions at 37·C and 5%$CO_2$. Cells were grown in DMEM (Gibco) supplemented with 10% FBS (PanBio), 1x Penicillin/Streptomycin, 1x Hepes and 1x Sodium Pyruvate.
***U2OS PHF13 Clone 5*:** U2OS cells were genetically modified with Tetracycline repressor and Tet-operated PHF13. This cell line was grown as described for U2OS cells, except that PHF13 was induced by the addition of Doxycycline (1ug/mL) to the medium. This cell line was verified for PHF13 expression using monoclonal antibodies against PHF13.
***293T*:** Human Embryonic Kidney cells (fetal) were grown under standard culture conditions at 37·C and 5%$CO_2$. Cells were grown in DMEM (Gibco) supplemented with 10% FBS (PanBio), 1x Penicillin/Streptomycin, 1x Hepes and 1x Sodium Pyruvate.

### METHOD DETAILS

**Cell lysis and Immunoblotting:** Fractionation of lysate was performed by lysing the pellet for 10 minutes on ice in CSK buffer (10mM PIPES, 100mM NaCl, 300mM Sucrose, 3mM $MgCl_2$, 0.1% NP40) supplemented with 1x complete protease inhibitor cocktail (Roche), followed by 5 minutes centrifugation at 2000g. Supernatant containing the cytoplasmic proteins was discarded and the pellet resuspended in mild chromatin buffer (100 mM NaCl, 20 mM Tris-HCl pH 7.5, 0.5% Triton-X, 2mM $CaCl_2$, 2.5 mM $MgCl_2$) plus complete protease inhibitors (Roche) for 10 minutes on ice, followed by 5 minutes centrifugation at 5000g. Supernatant was collected and stored as nucleoplasmic soluble fraction and the pellet was resuspended in mild chromatin buffer plus complete protease inhibitors and then sonicated on a Bioruptor Plus (Diagenode) for 10 cycles (30s on/ 30s off) at high intensity. Following sonication, 1 ul (250U) of Benzonase (Novagene) was added and RNA/DNA was allowed to degrade on ice for 30 minutes, followed by centrifugation at 14,000g for 5 minutes. Supernatant was collected (chromatin fraction) and any residual pellet was discarded. Total cell lysis was performed by lysing cells in RIPA buffer (50 mM Tris HCl pH 7.4, 150 mM NaCl, 1 mM EDTA, 1% NP40, 1% NaDoc, 0.1% SDS) supplemented with 1x complete protease inhibitor cocktail for 10 minutes on ice, followed by sonication on a Bioruptor Plus (Diagenode) for 10 cycles (30s on/ 30s off) at high intensity and then treatment with 1 µl (250U) of Benzonase (Novagene) on ice for 30 minutes, followed by centrifugation at 14,000g for 5 minutes. Supernatant (total cell lysis) was collected and residual pellet was discarded. Laemmli buffer was added and the lysates were denatured for 10 minutes at 99·C and then loaded on 4-15% precast polyacrylamide gels (Biorad) and transferred to PVDF membranes (Transblot Turbo-Biorad) using a Trans Blot Turbo Transfer System (Biorad) and the mixed molecular weight program. Membranes were then blocked for 2h at RT in 5% milk (in TBST) and then probed overnight with primary antibody. Membranes were then washed three times in 1x TBST followed by 1h incubation in secondary antibody in 5% milk (in TBST) at RT. Membranes were then washed three times in 1x TBST and then developed using Pico, Dura (Pierce) or Western Lightning Ultra (Perking Elmer) on a Biorad ChemiDoc XRS.

**Immunoprecipitation:** Immunoprecipitations were performed with $2x10^6$ cells per IP condition. Endogenous IP's were performed using 2 µg of specific antibody and 10 µl of magnetic protein A beads (DiaMag- Diagenode). IP's were either performed from chromatin fraction or RIPA lysates (see above) which were diluted 1:5 in dilution buffer (20mM Tris-HCl pH7.4, 100 mM NaCl, supplemented with 1x complete protease inhibitor and 2x PhosStop) or (50mM Tris-HCl pH 7.4, 150 mM NaCl, 1 mM EDTA supplemented with 1x complete protease inhibitor and 2x PhosStop), respectively, and then rotated overnight at 4°C. The next day the beads were washed 4x with the 1:5 diluted IP buffer (5 minutes each) and then resuspended in the lysis buffer supplemented with 1x Laemalli buffer, heated for 10 minutes at 99°C and loaded onto 4-15% gradient gels (Biorad). In the case of IP's performed from cells transfected with tagged proteins 25 µl of Flag-M2 agarose (Sigma) or 10 µl of magnetic GFP-Trap (Chromtek) were used instead of DiaMag Protein A beads.

**GST Protein production:** BL21 *ecoli* carrying pGEX4T3 -PHF13 Glutathione-S-transferase (GST) fusion proteins were grown in 50 mL LB-medium supplemented with 100 µg/mL ampicillin and 3.4 µg/ml chloramphenicol under rotation (200 rpm) at 35°C, overnight. 200 mL YT medium was inoculated with 5 mL of the overnight culture and incubated under rotation (200 rpm) at 37 °C until an $OD_{600}$ of 0.5 was reached. Expression of GST-fusion proteins was induced for 4 h at 32°C with 1 mM IPTG. After induction, the bacterial suspension was centrifuged for 15 min at 5000 rcf (4°C). Cell pellets were stored at -80°C overnight and then resuspended in 10 mL ice cold lysis buffer (50mM Tris-HCl pH8, 150 mM NaCl, 5mM EDTA, 1% Triton-X-100, 30 mg of Lysozyme and 1 complete protease tablet) and incubated for 10 min on ice. The lysate was then sonicated 5 times for 10 seconds and then centrifuged at 16,400 rcf for 30 min at 4°C. The clarified supernatant was then incubated with 500 µL of 50% glutathione agarose suspension in GST purification buffer (50mM Tris-HCl pH8, 100 mM NaCl, 1mM EDTA, 0.5% NP-40) and incubated under rotation for 2-4 h at 4°C. GST beads were then washed 4x in 10 mL GST purification buffer and briefly centrifuged (300 x g, 1 min, 4°C) in between. Finally, the beads were resuspended in 900 µL GST purification buffer and stored in aliquots at -80°C. The purity and quantity of prepared GST-fusion proteins bound to glutathione agarose was analyzed by SDS-PAGE.

**GST Pulldowns:** Cells were lysed on ice for 10 minutes with a CSK buffer (10mM PIPES, 100mM NaCl, 300mM sucrose, 3mM $MgCl_2$, 0.1% NP40, 1x complete protease inhibitor cocktail (Roche), 1 mM PMSF) and then centrifuged for 5 minutes a 2,000 x g. The cytoplasm was discarded and the pellet was then lysed for 10 minutes on ice in a mild chromatin buffer (20 mM Tris-HCl pH 7.5, 100 mM NaCl, 0.5% Triton-X 100, 2 mM $CaCl_2$, 2.5 mM $MgCl_2$ and 1x complete protease inhibitor cocktail (Roche)) to generate nucleoplasmic fractions. The cells were then centrifuged for 5 minutes at 10,000 x g. The supernatant was discarded and the pellet was then lysed in the mild chromatin buffer supplemented with 1x complete protease inhibitor cocktail (Roche) and benzonase (1 µl/3 x $10^6$ cells; Novagene 250U/ul 2698486) sheared for 10 minutes on a bioruptor (30s on 30s off at high intensity) and incubated on ice for 30 minutes. The cells were then centrifuged for 5 minutes at 10,000 x g and the chromatin supernatant was collected. GST pull downs were performed using 1 µg, 2.27 µg, 1.92 µg and 1.92 µg of recombinant GST, GST-PHF13, GST-ΔPHD and GST-ΔNTD proteins bound to Glutathione sepharose beads, respectively and nuclease digested chromatin lysate dilute 1:5 in dilution buffer (20 mM Tris-HCl pH 7.5, 100 mM NaCl and 1x complete protease inhibitor cocktail). The beads were incubated with the lysate at 4°C for 4h and then washed several times in the diluted chromatin buffer, prior to denaturation in Laemmli buffer and loading on a SDS-PAGE gel.

**In vitro translation:** In vitro translation was performed using a human cell free in vitro translation kit (Thermo Scientific-Cat.88855) according to the manufacturer’s instructions. Briefly, PHF13 and PHF13ΔNTD were cloned into the provided pT7-CFE1-CHis vector. A stop codon was inserted prior to the CHis to prevent the C-terminal fusion of PHF13 to the hexaHis tag. To obtain higher protein yields, the translation reaction was performed for 6-8 h and 2 h after the translation reaction was initiated additional 2 µL of transcription product were added. As a positive control, pT7-CFE1-CHis expressing eGFP was transcribed and translated in parallel and analyzed by fluorescence microscopy and Western blot. Expression of PHF13 and PHF13ΔNTD was verified by Western blot.

**Chromatography:** Chromatography was performed using recombinant proteins cleaved of their GST-tags (PreScission protease; GE Healthcare) on a Superose6 10/300 GL column (GE Healthcare) run in HMG250 buffer (25 mM HEPES (pH 7.5), 2.5 mM $MgCL_2$, 0.5 mM EDTA, 10% glycerol, 0,05% IGEPAL, 250 mM KCl) with a flow rate of 0.3 ml/min and a fraction size of 0.5 ml/min/fraction. Size exclusion chromatography of in vitro translated proteins (generated with a human cell free in vitro translation kit according to the manufacturer’s instructions; Thermo Scientific Cat:88855) was performed on a Superdex 200 5/150 GL column (GE Healthcare) run in 50 mM phosphate buffer (25 mM $KH_2PO_4$, 25 mM $Na_2HPO_4$ * $2H_2O$, 150 mM NaCl) with a flow rate of 0.3 ml/min/fraction.

**Cell Synchronization:** U2OS cells were grown to 70% confluence and then blocked with 2mM Thymidine for 20-24h. The cells were then released into fresh medium without Thymidine for 12h. 6h into the release, the cells were transfected with 10pM Control siRNA, PHF13 siRNA, SMC3 siRNA or PHF13si + SMC3si and 10 µl Lipofectamine 2000 (Thermo Fisher) according to manufactures instructions. Transfection reactions were left on the cells for 6h and then washed and split into several dishes and then placed into a second 1.25 mM Thymidine block for 18h to synchronize the cells at the G1/S border. The synchronized cells were then released in fresh medium for 7h allowing them to reach G2 and then blocked with 9 µM RO-3066 (Sigma; cdk1 inhibitor) for 6h to allow the cells to accumulate at the G2/M border. The cells were then released into mitosis by multiple washes with PBS, placed back in fresh medium and collected at 10-15 minute time intervals. For live cell analysis, SPY555 (Spirochrome) live cell DNA dye (0.5x staining solution according to manufactures instructions) was added to the medium with 10 µM verapamil for 2h before the final release from RO-3066 and was supplemented as well in the release medium to the same concentrations.

**Transfection:** Condensation experiments were performed in U2OS cells by transfection with Fugene 6 according to manufactures instructions using a 3:1 (reagent:DNA ratio) and 0.5-1 µg of PHF13 expression plasmid per 6-well for 24 h. siRNA experiments were performed by transfecting 10pM (control; 5’-AGGUAGUGUAAUCGCCUUG-3’, PHF13; 5’-UCACCUGUCCUGUGCGAAA-3’ or

SMC3; Origene SR306033) siRNA with 10ul Lipofectamine 2000 (ThermoFisher) according to manufactures instructions. Knockdowns were performed for 30-36h.

**FACS FRET:** FACS based FRET analysis were performed as previously described (Banning et al., 2010). In brief, 293T cells were transfected with 1µg of CFP and YFP constructs per 12-well using standard calcium-phosphate transfection for 24 hours. Cells were resuspended in FACS buffer (1x PBS, supplemented with 1% FBS and 0.5 mM EDTA) and double positive cells were analyzed for occurrence of FRET by fluorescence cell sorting with a BD-FACS Canto 2.

**Indirect Immunofluorescence:** Cells were grown on 10 mm round coverslips. Coverslips were fixed for 10 minutes with 4% paraformaldehyde at RT and then washed once with 1x PBS prior to permeabilization with 0.5% Triton-X for 10 minutes at RT, followed by 3 washes (5 minutes) with 1x PBS and 2h blocking in 5% BSA/PBS. Blocked coverslips were then incubated for 1h at RT with primary antibodies (using supplier recommended concentrations; or 1:200 PHF13 rabbit polyclonal antibodies or undiluted PHF13 rat monoclonal antibodies), followed by 3 washes (5 minutes) with 1x PBS and 1h incubation with secondary antibodies and DAPI or Hoeschst (Molecular Probes/Abcam 1:1000) and a final by 3 washes (5 minutes) with 1x PBS. Coverslips were then mounted on glass slides using Mowiol (Sigma) and allowed to harden before imaging. Pre-extraction IF experiments were performed using the same protocol except that the coverslips were first treated with CSK buffer (10mM PIPES, 100mM NaCl, 300mM Sucrose, 3mM $MgCl_2$, 0.1% NP40) for 20 minutes at RT and washed twice in PBS prior to 4% PFA fixation and 0.5% Triton-X permeabilization. Images were obtained by light microscopy using an Axiophot microscope (Zeiss), Observer (Zeiss) and Confocal microscope (Confocor 2; Zeiss).

**Microscopy**
To ensure, quantifiable linear signal values all image acquisitions followed strict rules to prevent over/under exposed pixels/used. All used signal intensities were adjusted by exposure time, dwell time and laser/led power to achieve at least 95 % of pixels within the lower 30 % of the maximal detectors digitalization ability. Furthermore, all images were digitalized in 8-bit. Mean Intensities are interpreted as densities, while sum intensities exhibit a total of stained structures/molecules. The standard deviation as factor for the homogeneity of a given signal.

**Live cell Imaging**
Living cells were imaged using the Cell Discoverer 7 Imaging Platform (Zeiss, Germany), in wide field mode running under ZEN Blue v 3.1 and full environmental control (5% v/v $CO_2$, 100 % humidity, 37 °C). Final experiments were performed using the Plan-Apochromat 20x / NA=0.95 objective, a 2x tubelens (Zeiss, Germany), and captured on an Axiocam 506 (Zeiss, Germany) with an 2x2 binning resulting in a lateral pixel resolution of 0.23 µm/ pix. Briefly, 5-10k U2OS cells were cultured and pretreated (knockdown, synchronization) in 96-well Screenstar plates (Greiner, Germany) or Ibidi 8 Well chambers (Ibidi, Germany). Living cells were stained in the microscope using Spy555-DNA (1:4000) and Spy650-TUB (1:2000) (Spirochrome, Germany) 2 h before synchronization release to allow cells and dyes to equilibrate and thermalize. To stabilize the Spy555-DNA nuclear signal we supplemented the medium with 10 µM verapamil simultaneously with the other dyes. The fully automated imaging approach captured 2-5 percent of individual well surfaces within 10 min intervals; focus stabilization was achieved by surface method, in the center of individual tile region, which typically consisted of 5x5 tiles and a total imaging area per well of 1.118 $mm^2$. All images were acquired with an additional transmitted light or contrasting method (Brightfield, Oblique, or Phase Gradient Contrast) channel.

**High Content Microscopy**
Fixed and immune fluorescence stained cells were imaged using the Celldiscoverer 7 Imaging Platform (Zeiss, Germany), in wide field mode running under ZEN Blue v 3.1 and temperature control (37 °C) to guaranty xyz-stage/sample stability over large scans. Again the Plan-Apochromat 20x / NA=0.7 objective (Zeiss, Germany) was used, followed by 2x tubelens (Zeiss, Germany), and camera binning of 3x3 binning resulting in a lateral pixel resolution of 0.347 µm/pix and a longitudinal resolution of 0.62 µm/pix. Briefly, 5-10k U2OS cells were cultured and pretreated (knockdown, synchronization,

block release, fixation and immune fluorescence staining) in 96-well Screenstar plates (Greiner, Germany). A total 40 % of the full well surfaces were imaged in 321 individual tiles per well. The full pipeline results typically in 395.472 single images, per experiment.

**Image Processing:** A dedicated ZEN v3.4 (Zeiss, Germany) stitched the live cell data based on nuclear counter staining as reference channels. For analysis in ImageJ the stitched, original images were exported without scaling or compression factors. High content 3D-image image data was first Maximum Intensity projected and subsequently stitched on the same workstation.

**Image Analysis: Zen -**Analysis of mitotic and dead indices in live cell experiments were performed using ZEN 3.4 Image analysis module. Briefly, stitched tiles from time-stacks were slightly smoothed (Gauss 1,3), intensity thresholded with fixed values to discriminate foreground and background, followed by water shedding. Dead, mitotic, and interphase cells were segregated by the following parameters listed in table.

| Cell State | Area ($\mu m^2$) | SD Nuclei (SPY555) | SD Nuclei (Bright field) |
|---|---|---|---|
| **Interphase** | >150 | < 6 | 0-15 |
| **Mitotitc** | >150 | > 6 | <15 |
| **Dead** | >100 | - | >15 |

Parameter overview cell state differentiation with ZEN 3.4 , Shown numbers were adjusted to the acquisition in 8-bit, and a pixel area of 0.051 $\mu m^2$.

**ImageJ:** Analysis of nuclear compaction was performed with customer Imagej and R scripts. Briefly, in a first attempt cells were picked from stitched time-stacks by identifying primary regions by mild smoothing, fixed intensity thresholds, close operation and water shedding. The resulting masks were filtered according to area within 100 $\mu m^2$-1000 $\mu m^2$ and circularity (sqrt((4*area)/($\pi$*FeretMax$^2$))) 0.4.-1.0. The resulting object was expanded by 200 pixels (~45 µm) and cropped out and saved. The Rois and resulting single cell time-stacks are stored in dedicated folders and used for quality experimenter quality control and further analysis.
A final script performed image analysis on single cell time stacks, by smoothing water shedding, filtering the resulting parameter tracks, as stated above.

**R:** A final customer R script row bound 500 individual tracks per condition and plotted the data in matrices using Topo. Colors as look up tables and stored data tables. All here stated scripts/snippets are public available https://github.com/ReneBuschow/SK_2021

**Live Cell Microscopy in Nikon Biostation:** Time-lapse microscopy was performed on a Nikon Biostation at 37°C and 5% $CO_2$**.** U2OS cells were transfected with 1 µg of PHF13 in 6cm dish and phase contrast images were collected every seven minutes over a period of 40 hours. Images were taken using a 40x objective at a resolution of 800 x 600 binning, exposure time of 1/40s and gain of 1.41.

**In Silico Data:** Ordered and putative interaction domains were predicted using Tango (http://tango.crg.es/). Disordered regions in PHF13 were called using PONDR (http://www.pondr.com/) (Obradovic et al., 2003), and inferred from AlphaFold2-advanced results. AlphaFold2-advanced (Mirdita M, Schütze K, Moriwaki Y, Heo L, Ovchinnikov S, Steinegger M. ColabFold - Making protein folding accessible to all. *bioRxiv*, 2021) was used to predict if PHF13 dimerizes and to identify interacting regions. The prediction of nucleolar localization was done using the NoD -nucleolar localization sequence detector (http://www.compbio.dundee.ac.uk/www-nod/) algorithm (Scott et al., 2010).

**Reprocessing of published ChIP-seq data:** For ChIP-seq analysis fastq files of PHF13, SMC3, RAD21, CTCF, H3K4me3, H3K27ac, and H3K9me3 were downloaded via Array Express (https://www.ebi.ac.uk/arrayexpress/ for PHF13) or fastq-dump (other experiments). Reads were mapped to mm10 reference genome using Bowtie2 (Langmead and Salzberg, 2012) (--very-sensitive), and if applicable, mapped reads from the same experiments but different sequencing runs were first merged and then filtered (-h -b -F 282 -q 10) with SAMtools (Li et al., 2009). Reads mapping to blacklisted regions (ENCFF547MET) were filtered out with SAMtools as well (Consortium, 2012). Signal tracks were computed with bamCoverage (-of bigwig -bs 10 -e 300 --normalizeUsing RPKM --ignoreDuplicates) (Ramirez et al., 2016). Regions with significant enrichment over input sample were identified using MACS2 (-g mm --keep-dup auto --bw 300 -q 0.05 -f BAM) (Zhang et al., 2008) using the --broad option for SMC3, RAD21, and PHF13 ChIP data.

| ChIP | ArrayExpress / GEO dataset | ID ChIP sample | ID control sample | doi |
|---|---|---|---|---|
| **PHF13** | E-MTAB-2636 | ERR689062 | ERR689061 | http://dx.doi.org/10.7554/eLife.10607.001 |
| **SMC3** | GSE80049 | GSM2111722, GSM2111723, GSM2111724 | GSM2111696, GSM2111697, GSM2111698 | https://doi.org/10.1038/s41588-017-0015-6 |
| **RAD21** | GSE74055 | GSM2099809 | GSM2099810 | https://doi.org/10.1038/cr.2018.1 |
| **CTCF** | GSE11431 | GSM288351 | GSM288358 | https://doi.org/10.1016/j.cell.2008.04.043 |
| **H3K4me3** | GSE120376 | GSM3399477 | GSM3399484 | https://doi.org/10.1186/s13059-019-1860-7 |
| **H3K27ac** | GSE120376 | | GSM3399484 | |
| **H3K9me3** | GSE120376 | | GSM3399484 | |

Heatmaps depicting ChIP-seq signal (RPKM normalized) distribution at selected genomic regions, were generated using computeMatrix (reference-point mode) and subsequently the plotHeatmap tool of the deepTools package (Ramirez et al., 2016).

**Correlation analyses:** Genome-wide correlation analyses for ChIP signal distributions were performed using multiBigwigSummary (deepTools,) (Ramirez et al., 2016) with RPKM normalized ChIP signals and a genome binning 2 kb bins (bins -bs 2000). The resulting count matrix of average scores per genomic bin was analyzed with plotCorrelation (deepTools) to obtain a clustered heatmap of pair-wise Pearson correlation coefficients.

**Molecular Dynamics (MD) simulations details:** The system composed by polymer beads and PHF13 molecules experiences thermal fluctuations at temperature T and the particles obey to the Langevin equation (Allen and Tildesley, 1989). For sake of simplicity, all monomers have same diameter $\sigma$ and mass m, both set to 1 in dimensionless units (Kremer and Grest, 1990). To account for excluded volume effects, we use between any two particles a repulsive Lennard-Jones (LJ) potential, with length scale $\sigma$ and energy scale $\varepsilon$ ($K_BT$ units). Consecutive beads of the polymer are linked by a finitely extensible non-linear elastic spring (FENE; (Kremer and Grest, 1990)), with standard parameter (Brackley et al., 2013; Chiariello et al., 2016) (length constant $R_0 = 1.6\sigma$ and spring constant $K_{FENE} = 30K_BT/\sigma^2$). Polymer stiffness is modelled through a standard three body interaction: $V_{stiff} = K_{stiff}(1 + \cos(\theta))$, where θ is the angle formed by three consecutive beads and $K_{stiff}$ is set to $2K_BT$. Bonds between monomers of PHF13 are modelled as harmonic springs: $V_{harm} = K_{harm}(r - r_0)^2$, with $r_0 = 1\sigma$ and spring constant $K_{harm} = 100K_BT/\sigma^2$).

All attractive interactions, i.e. protein-protein and protein-chromatin interactions, are modelled by a short-range, shifted attractive Lennard-Jones (LJ) potential $V_{LJ}$:

$$V_{LJ}(r) = 4\varepsilon\left[\left(\frac{\sigma}{r}\right)^{12} - \left(\frac{\sigma}{r}\right)^{6} - \left(\frac{\sigma}{R_{int}}\right)^{12} + \left(\frac{\sigma}{R_{int}}\right)^{6}\right]$$

for $r < R_{int}$, 0 otherwise. The interaction affinity $E_{A,B}$ between two generic types $A$ and $B$ is given by the minimum of $V_{LJ}(r)$ and is controlled by $\varepsilon$ and $R_{int}$. In our simulations, we consider specific affinities taken from the following ranges, which ensure the coil-globule transition of the polymer (Chiariello et al., 2016): $E_{PhD,H3K4me3} \in [6 \div 8]K_BT$, $E_{PhD,H3K4me1/2} \in [3 \div 4]K_BT$, $E_{NTD,NTD} \simeq 4K_BT$ and $E_{NLS,polymer} \in [2 \div 3]K_BT$. Simulations are performed with the software LAMMPS (Plimpton, 1995). The parameters defining the Langevin equation are set to standard values (Kremer and Grest, 1990): friction coefficient $\zeta = 0.5$, temperature $T = 1$ and integration time step $dt = 0.012$ (Rosa and Everaers, 2008), expressed in dimensionless units. The system is confined in a cubic simulation box with periodic boundary conditions, with edge size $D = 90\sigma$, in order to minimize finite size effects. Each simulation starts with the polymer initialized to a random Self-Avoiding-Walk (SAW) configuration (Chiariello et al., 2016) and PHF13 molecules uniformly distributed in the box, with a monomer concentration per volume unit $c = \frac{\left(\frac{4\pi r^3}{3}\right)N_{PHF13}}{D^3}$ sampled in the range $0.1\% \div 0.5\%$ , where $N_{PHF13}$ is the total number of PHF13 molecules and $r$ the radius of the monomer. For each parameter choice, we perform 10 independent simulations, which are equilibrated up to $3*10^7$ timesteps, so to ensure the coil-globule phase-transition. Polymer configurations are sampled every $10^6$ timesteps after the transition.

**Simulation of PHF13 mutations:** To simulate the effect of mutations on PHF13 molecules we act in general on the interaction potential between the PHF13 monomers involved in the mutation and chromatin binding sites, keeping the rest of the system mostly unchanged. Therefore, deletion of the PhD domain is modelled by silencing all the attractive interactions involving PhD, i.e. by setting $E_{PhD,H3K4me3} = E_{PhD,H3K4me1/2} = 0K_BT$ with $E_{NTD,NTD} \simeq 4K_BT$ and $E_{NLS,polymer} \simeq 3K_BT$.

Analogously, deletion of the NTD domain is modelled by silencing the attractive interactions involving NTD, i.e. by setting $E_{NTD,NTD} = 0K_BT$, with $E_{PhD,H3K4me3} = 6.1K_BT$ and $E_{PhD,H3K4me1/2} = 3.1K_BT$. Here, phase-separation of PHF13 is no longer observed, while residual interactions due to PhD-H3K4me3 persist. Upon reduction of $E_{PhD,H3K4me3}$ such interactions disappear, compaction is no longer observed and the polymer remains in an open, randomly folded configuration (Figure6).

**Polymer model including PHF13 and Cohesin:** Simulations including Cohesin are performed by introducing in the above-described system an additional type of molecule, modelled as a dimer of two beads having same diameter and connected by harmonic spring. Such molecules can bind to specific sites regularly located along the polymer (density ~0.2, two consecutive sites every 11 beads) with a very strong interaction ($8.1K_BT$), the other interaction affinities taken from the lower part of the above reported ranges. Concentration is taken as 10% of PHF13. Perturbations of the system are simulated as described before. Hence, depletion of Cohesin is simulated by simply switching off its attractive interactions and keeping unchanged the rest of the system. Conversely, depletion of the entire PHF13 is simulated by switching off all its attractive interactions with chromatin.

### QUANTIFICATION AND STATISTICAL ANALYSIS

### KEY RESOURCES TABLE

| Antibody Name | Clone Number | Company | Catalog |
|---|---|---|---|
| **Actin** | AC-15 | Sigma | A5441 |
| **Cyclin B** | Y106 | Epitomics | 1495-1 |
| **Flag** | M2 | Sigma | F1804 |
| **GAPDH** | 6C5 | Santa Cruz | SC-32233 |
| **GFP** | 7.1 and 13.1 | Roche | 11814460001 |
| **Histone H3** | E173-58 | Epitomics | 1326-1 |
| **Histone H3K4me3** | MC315 | Millipore | 04-745 |
| **Histone H3S10phos** | 6G3 | Cell Signalling | 9706 |
| **Ki67** | EPR3610 | Abcam | ab92742 |
| **NoH61** | | gift | |
| **Nucleolin** | MS-3 | Santa Cruz | Sc-8031 |

| | | | |
|---|---|---|---|
| **PHF13** | Rb polyclonal CR53 | gift Hans Will | |
| **PHF13** | Rb polyclonal S173 | gift Hans Will | |
| **PHF13 m33** | Rat monoclonal 7FB | Elizabeth Kremmer | |
| **PHF13 m34** | Rat monoclonal | Elizabeth Kremmer | |
| **PHF13 m45** | Rat monoclonal 6F6 | Elizabeth Kremmer | |
| **RAD21** | Rb polyclonal | Abcam | ab-992 |
| **RAD21** | B2 | Santa Cruz | SC-271601 |
| **SA2** | J-12 | Santa Cruz | SC-81852 |
| **SMC1** | EP2879Y | Abcam | ab75819 |
| **SMC3** | Rb polyclonal | Abcam | ab9263 |
| **SMC3-Ac** | 21A7 | Millipore | MABE1073 |
| **Topoisomerase 1** | Rb polyclonal | Biozol | Ab3825 |
| **Tubulin** | B-5-1-2 | Sigma | T5168 |

**Supplemental Information Titles and Legends:**

**Supplementary Figure 1: PHF13 is conserved and overlaps genome-wide with Cohesin and H3K4me3.** (A) Alignment of PHF13 proteins from evolutionary diverse species demonstrates the high conservation of the PHD domain (blue line) the NTD domain (green line), and the minimal homo-dimerization region (green dotted line). The PEST domains (black hatched line), NLS (yellow line) and Nucleolar localizing sequences (NoLS; light green dashed lines) are also depicted. (B) NoLS plot for PHF13 as predicted by NoD -nucleolar localization sequence detector algorithm. (C) Hierarchical clustering of genome-wide correlation (Pearson correlation) of ChIP-seq signals (RPKM normalized) with 2 kb binning. (D) Heatmap of PHF13, SMC3, and H3K4me3 ChIP-seq signal enrichment (RPKM) at PHF13 peaks, which do (upper panel) or do not (lower panel) overlap with SMC3 peaks. Each subset of PHF13 peaks was sorted by PHF13 signal enrichment.

**Supplementary Figure 2: Live and fixed cell analysis of PHF13 induced condensation** (A) 3D plot profiling H3S10 phosphorylation on the Z-axis in relation to the cell cycle plot in X and Y, demonstrating an overlap with the called mitotic cells and not the interphase cells, validating the gating called by SPY DNA staining. (B) Co-staining of DNA, PHF13 and H3S10 phosphorylation in U2OS cells throughout the cell cycle. (C) Visualization of mean PHF13 intensity over SD of DNA intensities, in different cell cycle phases, calculated from either standard or pre-extracted immunofluorescence protocols. Shown is a representative replicate out of 4 experiments. Data in this replicate is generated, gated, analyzed, from a total of 8961 pre-extracted and 17971 standard formaldehyde fixed single cells. (D) U2OS cells over expressing PHF13 that were Methanol-Acetone fixed and stained with PHF13 rat N- (4B2) and C- (7FB) terminal monoclonal antibodies. (E) Live-cell phase contrast microscopy images of U2OS cells transfected with PHF13 (1 µg) at different time points post PHF13 expression taken on a Nikon Biostation at 40x magnification. Red and blue arrows denote G1 or S nucleation, respectively. Imunofluorescence images were captured at 40x magnification on a Zeiss observer (B), a Zeiss Axiophot Phase Fluorescence microscope (D) and a Nikon Biostation (E).

**Supplementary Figure 3: Analysis of PHF13 oligomerization potential.** (A) Immunofluorescence images of U2OS cells transfected with CFP-fusion proteins as indicated that were used in FACS-FRET analysis. DNA was stained with Hoechst and images were captured on a Zeiss Axiophot Phase Fluorescence microscope. (B) Immunoblot analysis detected with a GFP antibody showing the expression of CFP- and YFP- PHF13 fusion proteins in transiently transfected 293 cells. Actin served as a loading control. (C) Immunodotblot of PHF13 full length and deletion mutants as separated by size exclusion chromatography (Superose6) and detected with a rat monoclonal 6F6 (PHF13 and ΔPHD) and 1D3 (ΔNTD and Δ24-40) PHF13 antibodies. (D) Immunoblot and Immunodotblot of *in vitro* translated full length PHF13 protein separated by size exclusion chromatography (Superdex 200) and detected with a rat monoclonal 1D3 anti PHF13 antibody.

**Supplementary Figure 4:PHF13 domain mutants.** (A) Schematic depiction of the PHF13 mutant proteins used to determine the essential features of condensation. (B) Immunoblot of PHF13 mutant proteins demonstrating similar expression level. Immunoblot was detectect with PHF13 rat monoclonal antibody 1D3. (C) Immunofluorescence images from U2OS cells transfected with PHF13 (1 µg) and stained for PHF13 (monoclonal antibody 6F6; red) and the nucleolar markers Topoisomerase I (green), NOH61 (green) and Nucleolin (green). DIC: Differential Interference Contrast. Images were captured on a Zeiss Confocor 2 confocal microscope.

**Supplementary Figure 5: PHF13 and Cohesin depletion impair mitotic condensation:** A) Live cell transmitted light images and Spy-550 DNA staining in green of the different siRNA knockdown conditions and control. B) Quantification of DNA compaction over time. Top: Two different look up tables of a mitotic cell. Depending on the DNA density, the color coding (Mean Fluorescence Spy-550-DNA staining) changes in the image. Bottom: Comparison of DNA densities of single cells released into mitosis. DNA densities of 500 cells each are plotted against time per condition (10 min intervals). C) Proportion of cell population H3S10 phosphorylation positive from fixed cells at different time points following release their release from R03066 under the specified experimental conditions (i.e. siCont., siPHF13, siSMC3 and siPHF13/siSMC3).

**Supplementary Figure 6: Molecular dynamics simulations**
A) Schematic of cohesin dimer, PHF13 molecule and chromatin bead polymer with corresponding distributions of binding sites. Cohesin is modelled as a dimer made of two identical beads, PHF13 as a molecule made of 3 beads representing N-terminal (NTD), NLS and PHD respectively. Chromatin filament is modelled as a standard chain of beads that includes binding sites representing H3K4me1/2 and H3K4me3 epigenetic marks and CTCF sites. B) Interaction affinities used to perform the simulations of the model including PHF13 and cohesin. C) Examples of 3D snapshots for the model with and without cohesin. D) Time dynamics of the model including only cohesion interactions.

**Supplementary Figure 7: Model predictions of PHD mutations are consistent with experiments**
A) PHD point mutations (PM) model. PHD is able to interact with cohesin while PHD-chromatin interactions are turned off. On the right, examples of three independent end-state structures. Upper panels: only the polymer is shown; bottom panels: cohesin, PHF13 molecules (red NTD domain) and polymer binding sites for cohesin are shown. White circles highlight PHF13 phase-separated clusters, induced by cohesin-PHD interaction. B) PHD deletion (**Δ**PHD) model. All interactions involving PHD are turned off. Examples of independent end-state structures. Note that no phase-separated PHF13 clusters are observed.
